# PI 4-kinases promote cell surface expansion and facilitate tissue morphogenesis during *Drosophila* cellularization and gastrulation

**DOI:** 10.1101/2022.09.09.507384

**Authors:** Wei Chen, Victoria Bergstein, Bing He

## Abstract

During epithelial morphogenesis, dynamic cell shape change driven by intrinsic or extrinsic forces requires prompt regulation of cell surface area. Using *Drosophila* ventral furrow formation as a model, we identified the PI 4-kinase Fwd as an important regulator for apical constriction-mediated cell shape changes. These morphological changes involve prompt lateral surface expansion in the constricting cells and apical surface expansion in the non-constricting cells adjacent to the constriction domain, both of which are impeded upon depletion of Fwd. Computer modeling demonstrates that restricting apical and lateral cell surface expansion will result in specific tissue-level morphological abnormalities during furrow formation, which well predicts the phenotypes observed in the *fwd* deficient embryos. Fwd also promotes cell surface expansion during cellularization, but this function is largely redundant with another PI 4-kinase, PI4KIIα. Together, our findings uncover an important role of Fwd in facilitating cell surface expansion in support of dynamic epithelial remodeling.

## Introduction

During tissue morphogenesis, cells frequently undergo dynamic shape changes in response to intrinsic or extrinsic cell-generated forces. One important task for cells during this process is to adjust their surface area to accommodate the rapid cell shape changes. Expanding surface area is particularly challenging as cell plasma membrane has low elasticity and cannot be extended by more than 3% to 5% without rupture ^1^. Despite this challenge, many studies have shown that under physiological conditions, cells can manage to expand their surface area in face of the mechanical stress and still maintain membrane integrity ^2–5^. The ability of cells to promptly alter their surface area under mechanical stress brings several central questions: how do cells adjust surface area while rapidly changing their shape? If active processes are involved, how do they coordinate with cell shape change? What are the consequences if cells fail to adjust surface area? Because cell shape changes are one of the fundamental cellular processes underlying tissue morphogenesis, answers to these questions are essential for understanding the mechanisms of tissue construction in development.

Ventral furrow formation during *Drosophila* gastrulation provides a good model to address those questions. The morphogenetic changes of ventral furrow formation have been well documented ^6, 7^. Before ventral furrow formation, embryo undergoes a special cleavage called cellularization. Cellularization begins with a syncytium with approximately 6000 nuclei aligned at the periphery of the embryo. During cellularization, plasma membrane invaginates to form cleavage furrows between the neighboring nuclei and separate them into a monolayer of epithelial cells ^8^. At the end of cellularization, a group of ventrally localized mesodermal precursor cells, determined by Dorsal nuclear gradient and its downstream transcription factor Twist and Snail ^9, 10^, undergo apical constriction and invaginate into a ventral furrow. Ventral furrow formation proceeds in two steps ^7^. First, apical constriction induces a 70% increase in apical-basal cell lengthening (“lengthening phase”). This is then followed by cell shortening and internalization as the tissue invaginates (“shortening phase”). The whole process lasts about twenty minutes and results in the formation of an anterior-posteriorly oriented furrow. The molecular mechanism of apical constriction has been well characterized. The expression of Twist and Snail in the mesoderm precursor cells results in the activation of the Fog-G protein coupled receptor (GPCR) pathway, which triggers the downstream RhoA/Rho1 signaling cascade. The Rho1 signaling leads to apical assembly of F-actin and activation of non-muscle myosin II (henceforth “Myosin II”), which form an actomyosin network that drives apical constriction through pulsed contractions (reviewed in ^11–13^).

The mechanism that mediates cell lengthening during the early phase of ventral furrow formation has been recently elucidated. Quantitative analysis of 3D cell shape dynamics revealed an intimate coupling between apical constriction and cell lengthening ^14^, which led to the proposal that gastrulating cells lengthen via a purely mechanical mechanism. The cytosol acts as an incompressible fluid, and constriction of apical actomyosin network generates forces that drive basally directed movement of the cytoplasm, resulting in cell lengthening ^14^. In line with this view, a later study demonstrates that during apical constriction, the tissue interior behaves like a viscous continuum and undergoes a tissue-scale laminar flow, which causes apical-basal lengthening of the constricting cells ^15^. Interestingly, during this process, the lateral membrane of the cells appears to readily expand with the flow without impeding the flow, suggesting a close coupling between cell surface expansion and the viscous movement of the cytoplasm ^15^. This coupling is unlikely achieved through spreading of preexisting membrane folds, as previous EM studies show no obvious membrane folding prior to ventral furrow formation ^16–18^. Thus, apical constriction-induced cell lengthening likely involves mechanisms that add new membrane material to the cell surface.

In this study, we identified Four-wheel drive (Fwd) as an important factor regulating cell surface expansion during the lengthening phase of ventral furrow formation. Fwd is a *Drosophila* homologue of mammalian PI 4-kinase, PI4KIIIβ. Four members of PI 4-kinases have been identified in mammals: PI4KIIα, PI4KIIβ, PI4KIIIα, PI4KIIIβ ^19^. *Drosophila* genome encodes three PI 4-kinases corresponding to the mammalian homologues except for PI4KIIβ ^20^. PI 4-kinases catalyze the production of PI(4)P by phosphorylation of phosphatidylinositol (PtdIns) ^19, 21^. Previous studies have revealed various important roles for PI(4)P in regulating the endomembrane system. PI(4)P regulates Golgi-to-plasma membrane transport by facilitating vesicle formation from trans-Golgi-network (TGN) ^22, 23^. PI(4)P also serves as a recognition site for lipid transport protein such as OSBPs, FAPPs and CERTs, which mediate lipid transport from ER to the Golgi ^24^. Finally, PI(4)P serves as a precursor for other important phosphoinositide species, such as PI(4,5)P_2_ ^25, 26^. Different PI 4-kinase has distinct subcellular localization and participates in different cellular processes ^19, 21^. PI4KIIIβ localizes to TGN, where it functions partially redundantly with PI4KIIα, another Golgi-localized PI4 kinase, in promoting antegrade trafficking from TGN ^19, 27^. In addition, PI4KIIIβ itself can serve as scaffolding protein to recruit Rab11 to facilitate vesicle trafficking from TGN ^28–30^. During *Drosophila* spermatogenesis, both kinase and scaffolding activities of Fwd have been demonstrated to facilitate vesicle trafficking that are important for successful meiotic cytokinesis ^31^.

Despite the important function of PI4KIIIβ in regulating Golgi-to-plasma membrane transport, null mutant flies are viable. The only reported phenotypes in *Drosophila* are cytokinesis defect during spermatogenesis and mitochondria dysfunction in neuron and muscle cells ^4, 31–33^. In this study, we uncover a previously unappreciated role of Fwd in regulating cell surface area during early embryogenesis. During ventral furrow formation, Fwd facilitates furrow invagination through its separate functions in promoting cell surface expansion and regulating apical Myosin II network organization. During cellularization, Fwd functions redundantly with PI4KIIα to promote cleavage furrow ingression. Together, these findings demonstrated an important role for PI 4-kinases in promoting cell surface expansion during epithelial morphogenesis.

## Results

### Cell surface area increases during apical constriction-mediated cell lengthening

During the lengthening phase of ventral furrow formation, cells undergoing apical constriction elongate along the apical-basal axis by a factor of ∼1.7 (Figure 1A) ^7^. The change of cell morphology from columnar to a more slim and elongated shape predicts a significant increase in the cell surface area (Figure 1B). To test this prediction, we imaged gastrulating embryos expressing Ecadherin-GFP and segmented individual cells in 3D following a recently described protocol ^34^ (Figure 1C). Consistent with the prediction, we observed an increase in both cell length and cell surface area during apical constriction (Figure 1D, E). The average rate of cell surface area increase was 28.5 μm^2^/min, which resulted in an average of 26.0% ± 8.2% (mean ± s.d, n = 15 cells from 3 embryos) increase of surface area within the first 8 minutes of ventral furrow formation (Figure 1D). Accordingly, the average cell length increased by 53.0% ± 4.2% (mean ± s.d, n = 15 cells from 3 embryos) within the same time span at an average rate of 2.6 μm/min (Figure 1E). Our measurement thus confirmed that apical constriction induced cell lengthening involves a net increase of cell surface area.

**Figure 1.**
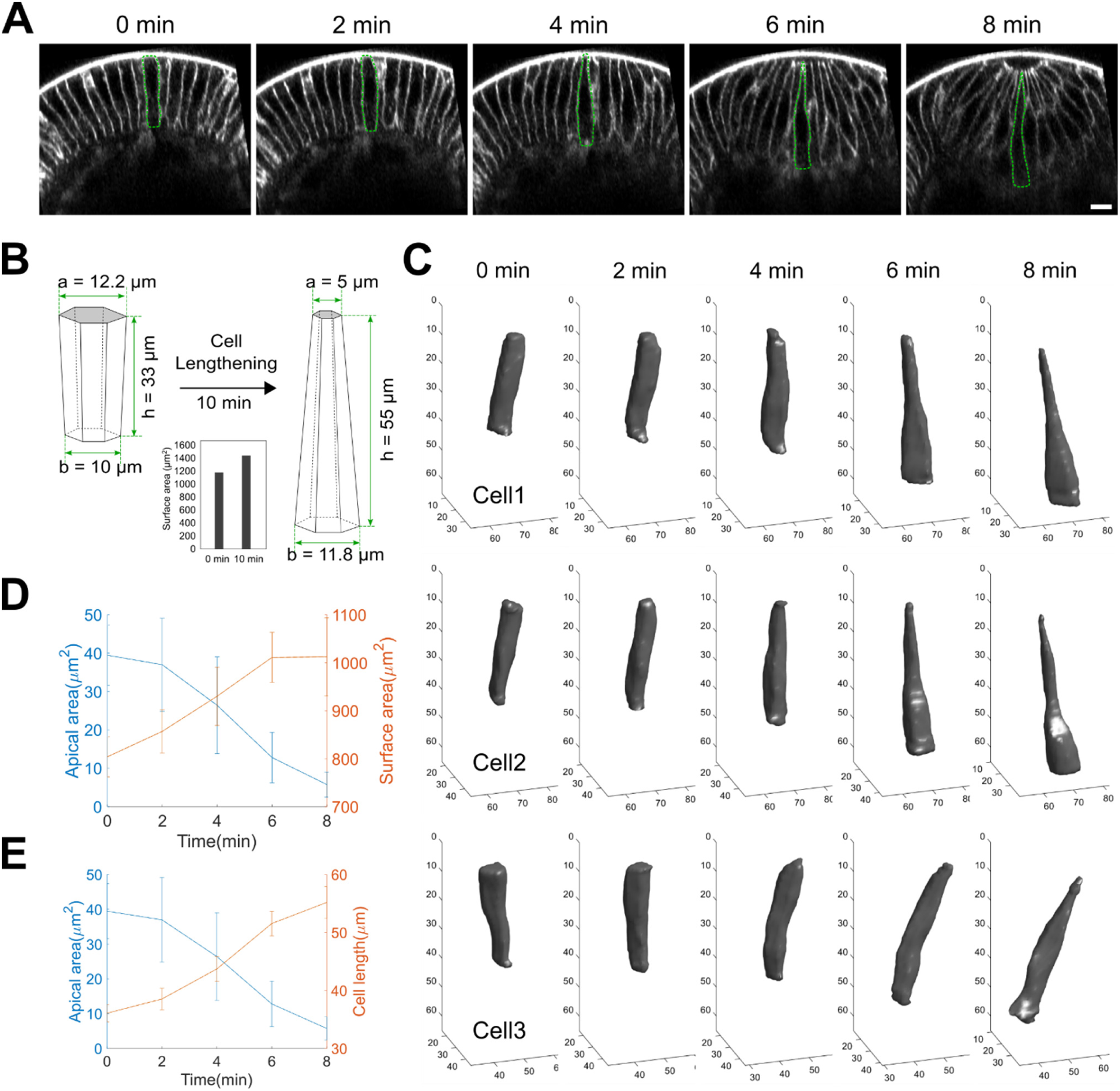
Cell surface expands during apical constriction induced cell lengthening. **(A)** Cross-section view of an embryo expressing E-cadherin-GFP showing ventral furrow formation. Ventral side is up. For each frame, the outline of a single cell is shown as an example. Scale bar: 10 μm. **(B)** Predicted cell surface area increase based on the observed cell shape change in the constriction domain. **(C)** Examples of three-dimensional reconstruction of ventral cells from a wildtype embryo during apical constriction. **(D, E)** Apical area and cell length (D) or cell surface area (E) over time during apical constriction. N=15 cells from 3 embryos. Error bars stand for s.d.

### Disruption of *Drosophila* PI4KIIIβ homologue Fwd results in reduced rate of cell lengthening during apical constriction

Previous electron microscopy studies demonstrate that no obvious plasma membrane infoldings are present in the lateral membranes of the cellular blastoderm prior to gastrulation^17, 18^. In addition, apical cell microvilli, which serve as a membrane reservoir for membrane ingression during cellularization, have been depleted at the end of the cellularization ^16–18, 35^. Therefore, it is unlikely that cell surface expansion during apical constriction is achieved through unfolding of pre-existing membrane folds. Given the non-stretchable nature of the plasma membrane ^1^, we hypothesized that cell lengthening/surface area increase requires membrane supply from intracellular pool of membrane. Exocytic membrane trafficking is a tempting mechanism in this process since it has been shown to promote surface area increase in many other cell shape change related processes ^36^.

To investigate the molecular mechanism of cell surface expansion in cells undergoing apical constriction, we performed a candidate RNAi screen for genes involved in exocytic trafficking to search for mutants with defective cell lengthening during ventral furrow formation (Methods; Table 1). For the majority of the candidates, we observed mild or no obvious cell lengthening defects and embryos appeared to develop normally through early gastrulation (Table 1). This might be due to either low RNAi efficiency or a non-essential function of the candidate gene during the stage of interest. A subset of candidate RNAi showed severe early defects such as lack of egg laying or abnormalities beyond cell lengthening defect (Arf1, Sec5, Rab11, dynein, PI4KIIIα), indicating that the target genes are essential for other developmental processes earlier than gastrulation. However, from this screen, we identified one interesting candidate, Fwd, the *Drosophila* homologue of PI4KIIIβ that is important for the Golgi to plasma membrane trafficking. RNAi mediated maternal knockdown of *fwd* resulted in reduced cell lengthening during apical constriction (Figure 2A-E). In the control wildtype embryos, cell lengthening proceeded in two consecutive phases. In the first 3-4 minutes of apical constriction (T = 0 indicates the onset of apical constriction, same below), cell length increased at an average speed of 1.6 μm/min (Figure 2B-E, cyan box). At around T = 4 minutes, cell lengthening speeded up and proceeded at an average speed of 2.6 μm/min (Figure 2B-E, magenta box). In the *fwd* RNAi embryos, cell lengthening proceeded at an average speed of 1.7 μm/min during phase 1, which was comparable to the wildtype embryos. However, in phase 2, the average speed in the *fwd* RNAi embryos reduced to 1.1 μm/min (Figure 2B-E). In addition, it took longer time for *fwd* RNAi embryos to reach the maximum cell length. In the control embryos, cell length increased from 35 μm (35.4 ± 1.5, mean ± s.d.) at the onset of apical constriction to a maximum of 55 μm (55.0± 1.7, mean ± s.d.) in approximately 10 minutes. In *fwd* RNAi embryos, cell length increased from 29 μm (29.3 ± 1.1, mean ± s.d.) to a maximum of 48 μm (48.2 ± 1.9, mean ± s.d.) in approximately 14 minutes (Figure 2C). These results demonstrate that the rate of cell lengthening during apical constriction is substantially reduced in *fwd* RNAi embryos. We observed similar cell lengthening defects in embryos derived from *fwd*^3^/Df females ^32^ (Figure 2A, B), suggesting that the lengthening phenotypes in the *fwd* RNAi embryos are unlikely due to off-target effect of RNAi.

**Figure 2.**
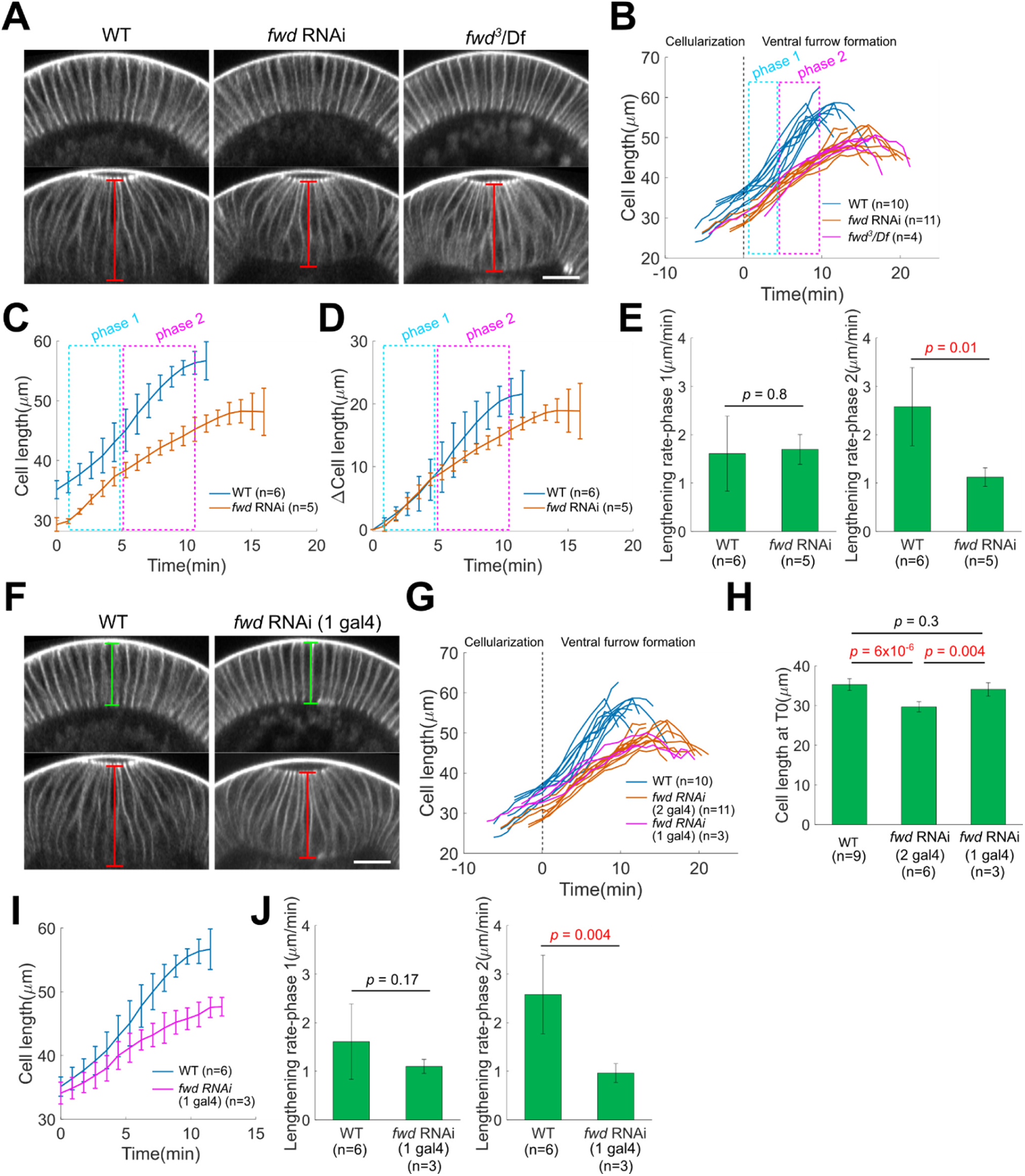
Fwd knockdown results in cell lengthening defect during ventral furrow formation. **(A)** Representative cross-section views showing wildtype, *fwd* RNAi and *fwd* loss of function mutant embryos at the onset of ventral furrow formation (top) and near lengthening-shortening transition (bottom). Apical-basal length of the constricting cell at the center of the furrow (red lines) was measured over time. Scale bar, 20 μm. **(B)** Cell length over time for different genotypes. In wildtype embryos, cell lengthening proceeds in two phases with distinct rates (cyan and magenta boxes). *fwd* loss of function mutants and *fwd* RNAi embryos exhibit similar lengthening defect. 0 min is defined as the onset of apical constriction throughout the text. **(C, D)** Average cell length over time before (C) or after (D) subtracting the cell length at 0 min. Error bars stand for s.d.. **(E)** Rate of cell lengthening at phase 1 and phase 2. Error bar stands for s.d.. **(F)** Representative cross-section views showing wildtype embryo and *fwd* RNAi embryo with one copy of Gal4. Top: onset of ventral furrow formation. Bottom: near the lengthening-shortening transition. Green and red lines indicate cell length at different stages. Scale bar, 20 μm. **(G)** Cell length change over time. For G and H, Data for WT and *fwd* RNAi (2 gal4) are reused from (B). **(H)** Cell length at the end of cellularization. Error bar stands for s.d.. For H – J, Data for WT are reused from (B). **(I)** Average cell length over time. Error bars stand for s.d.. **(J)** Rate of cell lengthening at phase 1 and phase 2. Error bar stands for s.d.. Two tailed, unpaired Student’s t test is used for all statistical analysis shown in this figure.

**Table 1:** Candidate RNAi and dominant negative lines used for the screen for lengthening defects and the phenotypes in ventral furrow formation.

| Gene | Inhibition method | Bloomington Stock Number | Ventral furrow formation phenotype |
| --- | --- | --- | --- |
| Rab4 | RNAi | BL33757 | Normal |
| Rab8 | dominant negative | BL9780 | Abnormal cell shape |
| Rab11 | dominant negative | BL9792 | No egg laying |
| Arf1 | RNAi | BL66174, BL66175 | Mild lengthening defect and abnormal cell shape |
| Sec71 | RNAi | BL50539 | Normal |
| Garz | RNAi | BL34987 | Mild lengthening defect |
| Nuf | RNAi | BL44035 | Normal |
| Crag | RNAi | BL53261 | Normal |
| Brun | RNAi | BL64934 | Normal |
| Gga | RNAi | BL36905, BL51170 | Normal |
| dynein | RNAi | BL36698 | No egg laying |
| dynein | RNAi | BL36583 | Mild lengthening defects |
| MyosinV | RNAi | BL55174 | Normal |
| Cert | RNAi | BL35579, BL60080 | Mild lengthening defect |
| PI4KIIα | RNAi | BL35278, BL65110 | Mild lengthening defects |
| PI4KIIIα | RNAi | BL35643 | No egg laying |
| Fwd | RNAi | BL35257 | Lengthening defects |
| Sac1 | RNAi | BL56013 | Normal |
| Sec5 | RNAi | BL50556 | Mild lengthening defect and abnormal cell shape |

Note: Phenotypes were assessed based on visual inspection.

We noticed that the cells in *fwd* RNAi embryos at the onset of cellularization was 17% shorter than that in the control embryos, suggesting that Fwd also plays a role in cell growth during cellularization (Figure 2C). When we lowered the expression level of *fwd* shRNA by reducing the copy number of the *GAL4* driver, the ventral cell length at the onset of apical constriction became comparable to that in the wildtype embryos, yet cell lengthening during apical constriction was still affected to a similar extent as in the original *fwd* knockdown embryos (Figure 2F-J). This result demonstrates that the cell lengthening defects in *fwd* RNAi embryos are not due to the defect in the starting cell length at the onset of gastrulation.

### Depletion of *fwd* leads to mild defects in apical constriction, but this defect cannot fully account for the defects in cell lengthening

Previous work demonstrates that apical constriction and cell lengthening are tightly coupled ^14, 15^. It is therefore possible that the cell lengthening defects we observed in *fwd* RNAi embryos were a secondary consequence of the apical constriction defects. To test this possibility, we quantified the rate of apical constriction in control and *fwd* RNAi embryos by measuring the total length of the apical surface of the constricting cells in the 2D cross-section view (the “apical width”, red lines in Figure 3A; Methods). The decrease of apical width over time was largely comparable between wildtype and *fwd* RNAi embryos, except that the rate was slightly lower in *fwd* RNAi embryos (Figure 3B, C). Thus, depletion of Fwd leads to a moderate but detectable reduction in the rate of apical constriction.

**Figure 3.**
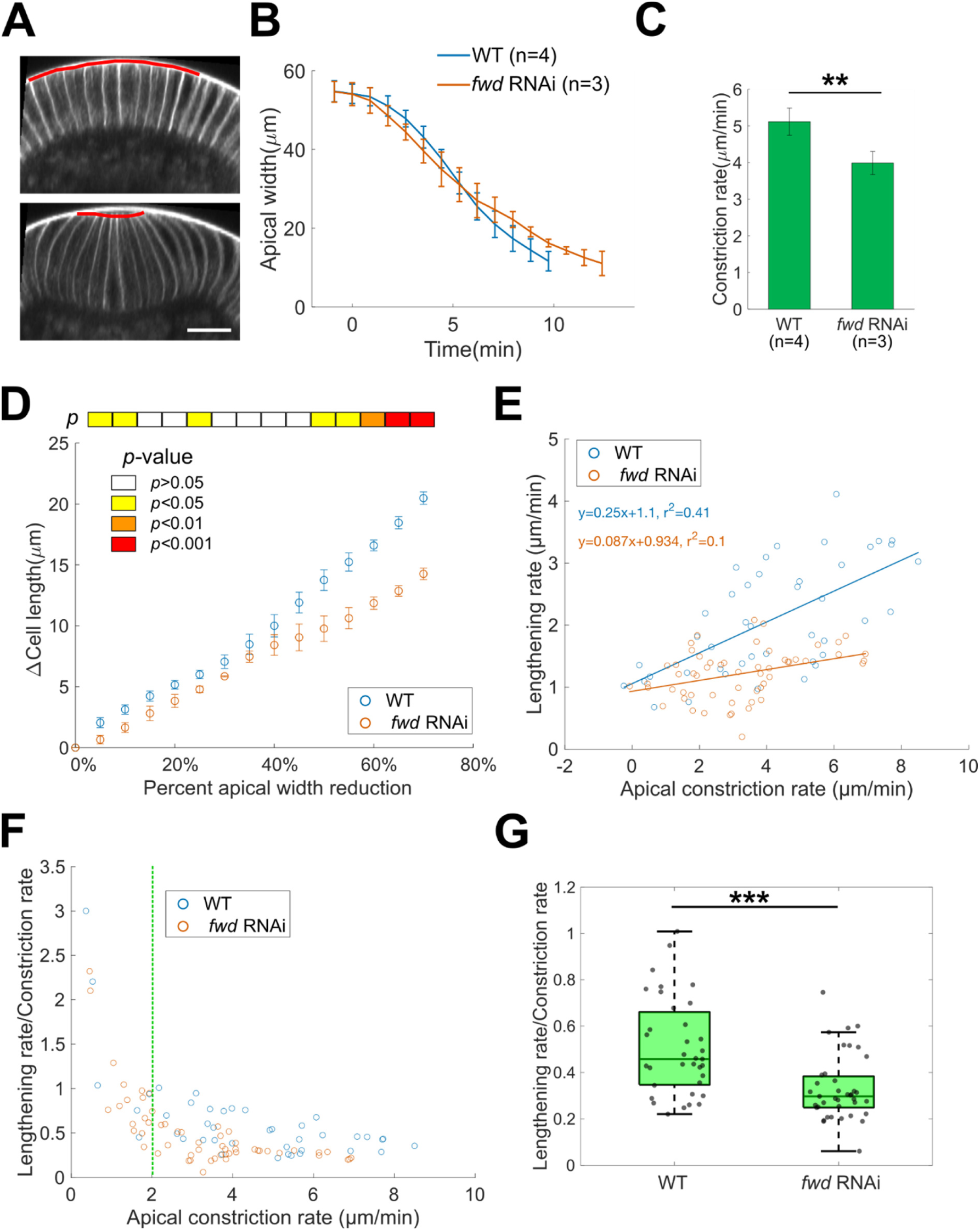
Knockdown of *fwd* leads to mild defects in apical constriction, but this defect cannot fully account for the defects in cell lengthening. **(A)** Example showing the measurement of apical constriction domain width from the cross-section views (red lines). A row of ∼10 cells centering around ventral midline were measured over time (Methods). **(B)** Average apical constriction domain width over time. Error bars stand for s.d.. **(C)** Constriction rate during 3 – 10 min. **: p < 0.01. Two tailed, unpaired Student’s t test. **(D)** Cell length increase as a function of percentage apical area reduction. With the same degree of apical constriction, *fwd* RNAi embryos show less cell lengthening compared to wildtype embryos. Error bars stand for s.e.m.. p-values are shown as a color bar on the top of the plot. Two tailed, unpaired Student’s t test. **(E)** Scatter plot of cell lengthening rate as a function of constriction rate. **(F)** Scatter plot of the ratio between cell lengthening rate and apical constriction rate as a function of apical constriction rate. For the analysis shown in (G), data points with constriction rate smaller than 2 μm/min (green dotted line) were excluded. **(G)** Comparison of the cell lengthening rate/apical constriction rate ratio between wildtype and *fwd* RNAi embryos. ***: p < 0.001. Two tailed, unpaired Student’s t test.

Next, we asked whether the difference in apical constriction rate could account for the difference in the rate of cell lengthening between wildtype and *fwd* RNAi embryos. We reasoned that if the lengthening defect in *fwd* RNAi embryos was entirely caused by the reduction in the rate of apical constriction, the relationship between apical constriction and cell lengthening should be the same as in the control embryos. To test it, we compared the extent of cell lengthening between the two genotypes when the same extent of apical constriction was achieved. In control embryos, the increase in cell length was quasi-linearly related to percent apical width reduction (Figure 3D). In *fwd* RNAi embryos, the increase in cell length upon the same level of apical constriction was consistently lower compared to the control embryos, and the difference between the two genotypes became more prominent as apical width reduction exceeds 50% (Figure 3D).

To further analyze the relationship between apical constriction and cell lengthening, we examined the rate of cell lengthening during every one-minute interval as a function of the rate of apical width reduction during the same time span. In the control embryos, there was a moderate positive correlation between the two rates (r^2^ = 0.4; Figure 3E). Knockdown of *fwd* resulted in reduced lengthening rates compared to the control at comparable constriction rates (Figure 3E). As a result, the correlation between the constriction rate and the lengthening rate became negligible for *fwd* RNAi embryos (r^2^ = 0.1; Figure 3E). In support of this observation, the ratio between cell lengthening rate and constriction rate was significantly higher in wildtype embryos compared to *fwd* RNAi embryos (Figure 3F, G). Together, these results indicate that although *fwd* knockdown had a moderate effect on the rate of apical constriction, this effect could not fully account for the reduced rate of cell lengthening in the knockdown embryos.

### The cell lengthening defect in *fwd* knockdown embryos is associated with defect in cell surface expansion

To determine whether the cell lengthening defect in *fwd* RNAi embryos reflects defect in cell surface area increase, we performed 3D cell segmentation for the constricting cells in *fwd* RNAi embryos (Figure 4A). In support of our hypothesis, we found that depletion of Fwd indeed resulted in prominent defects in cell surface expansion during apical constriction, as indicated by a reduced rate of cell surface area increase (control: 0-8 min: 28.5 ± 6.7 μm^2^/min; *fwd* RNAi: 0-12 min: 11.6 ± 3.5 μm^2^/min) (Figure 4B-C). As a result, the overall amount of cell surface area increase during the lengthening phase was substantially reduced in *fwd* RNAi embryos (control: 208 ± 65 μm^2^ over 8 minutes; *fwd* RNAi: 127 ± 47 μm^2^ over 12 minutes) (Figure 4B-C). In addition, the increase in apical-basal cell length was also reduced in *fwd* RNAi embryos, in agreement with the 2D measurement (Figure 4D-E).

**Figure 4.**
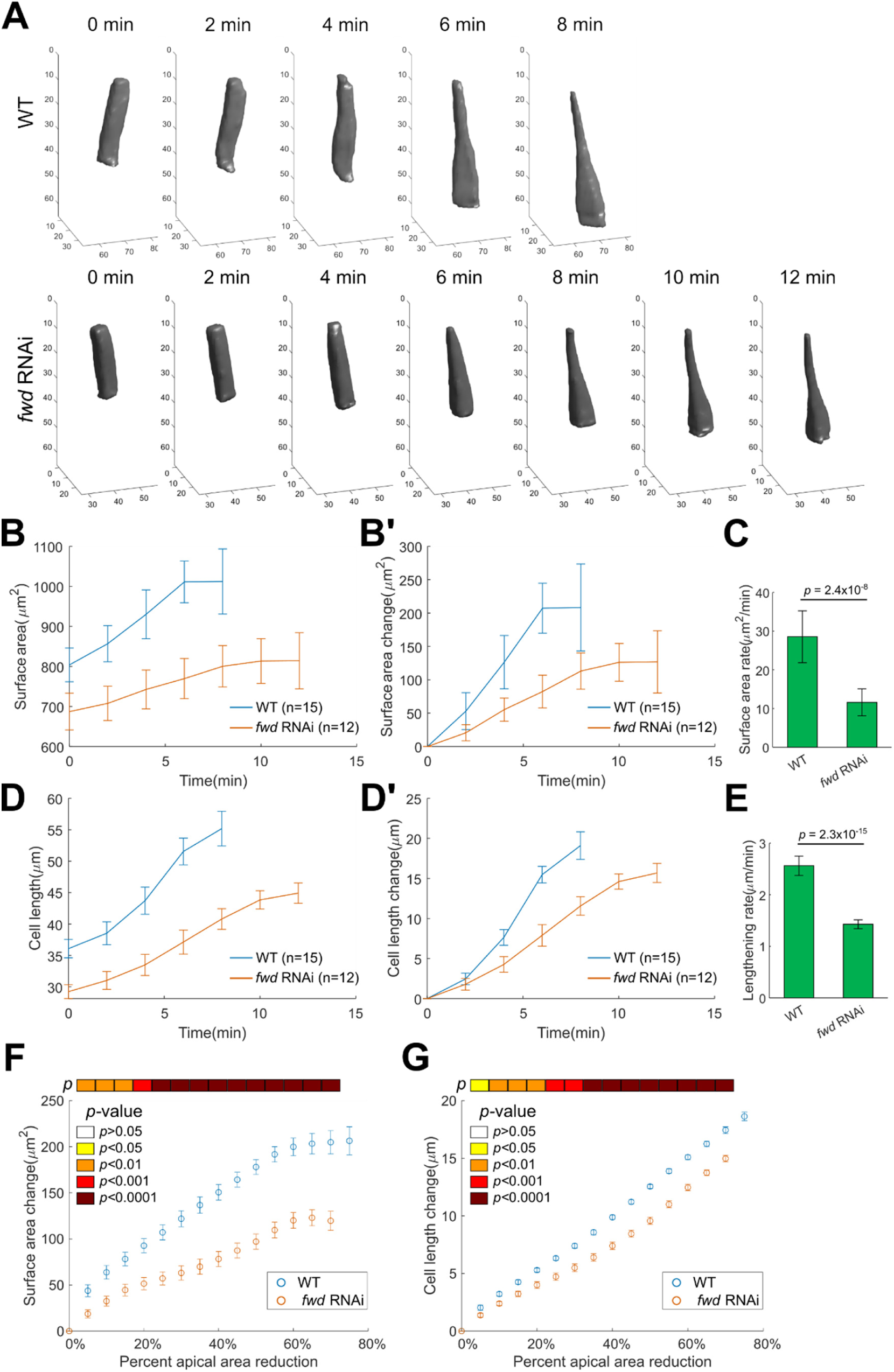
Cell lengthening defect in *fwd* RNAi embryos is associated with reduced cell surface expansion. **(A)** Representative three-dimensional reconstruction of ventral cells undergoing apical constriction in wildtype and *fwd* RNAi embryos. Image sequence begins at the onset of gastrulation and ends ∼ 2 min before lengthening-shortening transition. The wildtype data shown in Figure 1 were reused in this figure for comparison. **(B, B’)** Average cell surface area over time before (B) or after (B’) subtracting the value at 0 min. For wildtype, N=15 cells from 3 embryos; For *fwd* RNAi, N=12 cells from 3 embryos (same below). Error bars stand for s.d.. **(C)** Rate of the surface area change during the lengthening phase. Error bars stand for s.d.. **(D, D’)** Average cell length over time before (D) or after (D’) subtracting the value at 0 min. Error bars stand for s.d.. **(E)** Rate of the cell length change during the lengthening phase. Error bars stand for s.d.. **(F, G)** Cell surface area increase (F) and cell length increase (G) measured from 3D reconstructed cells as a function of the percentage decrease of apical area. The percentage decrease of apical area was measured by averaging the apical area of 80 wildtype and 72 *fwd* RNAi cells located at the mid-ventral region of the embryo (from 3 wildtype and 3 *fwd* RNAi embryos, respectively). Error bars stand for s.e.m.. p-values are shown as a color bar on the top of the plot. Two tailed, unpaired Student’s t test is used for all statistical analysis in this figure.

Of note, the cell length and cell surface area in *fwd* RNAi embryos were 19.0% and 14.5% smaller than the control embryos at the onset of gastrulation (Figure 4B, D; Cell length: control: 36.1 ± 1.5 μm, *fwd RNAi*: 29.2 ± 1.1 μm; Cell surface area: control: 804 ± 42 μm^2^, *fwd* RNAi: 687 ± 46 μm^2^), consistent with the 2D measurement. In addition, the 3D analysis also confirmed the reduced rate of apical constriction in *fwd* RNAi embryos (Supplementary Figure 1A). Using a similar strategy as described above, we show that the difference in the rate of apical constriction could not fully account for the reduced rate of cell surface expansion and cell lengthening in *fwd* knockdown embryos (Figure 4F, G).

Interestingly, we noticed a difference in cell volume change between the control and *fwd* RNAi embryos. In the control embryos, there was a moderate, 12% ± 11% increase in cell volume within the first 8 minutes of apical constriction (Supplementary Figure 1B). A small increase in cell volume during the first few minutes of apical constriction has been previously reported ^14^. In *fwd* RNAi embryos, however, no overall volume increase was observed within the first 12 minutes of apical constriction (Supplementary Figure 1B). The cause of the cell volume phenotype in *fwd* RNAi embryos and its link with the cell surface area phenotype remain to be determined.

Together, our observations demonstrate that Fwd is required for proper cell surface area increase during the lengthening phase of ventral furrow formation. While Fwd also plays a role in promoting apical constriction, our analysis suggests that Fwd regulates cell lengthening and cell surface expansion through mechanisms that are distinct from its function in apical constriction.

### Fwd regulates the coordination of apical area reduction and the spatial organization of apical Myosin II in the constricting cells

The apical constriction defects observed in *fwd* RNAi embryos prompted us to examine whether Fwd also regulates cell surface area on the apical side of the cells. In addition to the mild reduction in apical constriction rate, we noticed another interesting phenotype, that the apical domain of the constricting cells appeared more uniform in *fwd* RNAi embryos compared to the control embryos. To further analyze this phenotype, we generated the flattened surface view of the embryo (Methods) and analyzed apical cell morphology at a defined stage when apical constriction reaches close to 50% (Figure 5A) ^37^. Specifically, we focused on the distribution of apical cell area and anisotropy across the central region of the constriction domain (∼6-7 cells spanning across the ventral midline, Figure 5B). As expected, the average cell area was similar between the two genotypes, which validated the selection of stage for the analysis (Figure 5C). The average cell anisotropy was also similar between the control and *fwd* RNAi embryos (Figure 5D). Despite these similarities, the constricting cells in the *fwd* RNAi embryos appeared to be much more homogeneous in size and shape (Figure 5B), as reflected by the significantly smaller variations in both cell area and cell anisotropy (Figure 5E, F). This difference only emerged during apical constriction. Prior to gastrulation, the prospective constricting cells appeared homogeneous in both wildtype and *fwd* RNAi embryos (Figure 5G-K).

**Figure 5.**
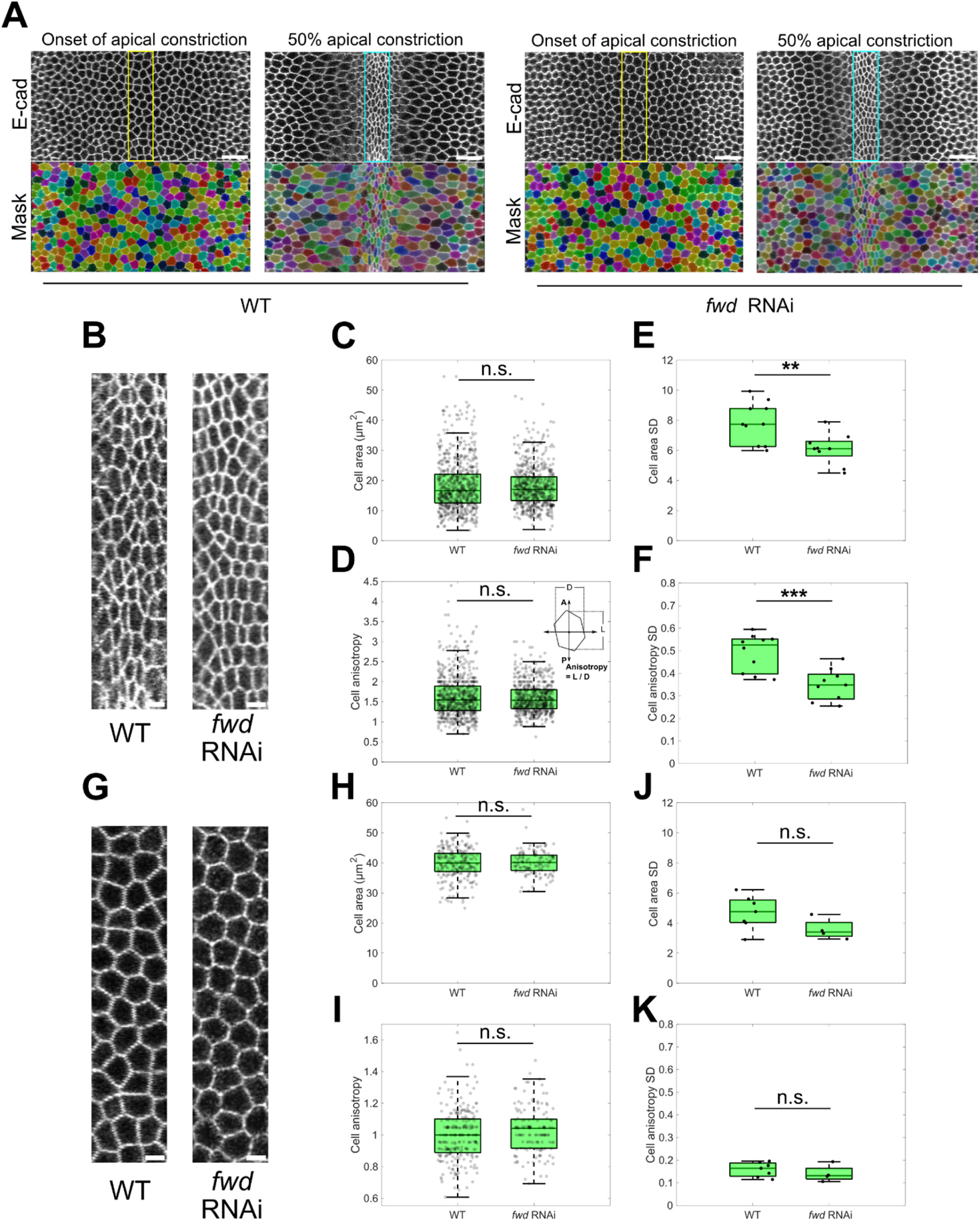
Depletion of Fwd results in more uniform cell shape in the constriction domain. **(A)** Cell segmentation at the ventral surface of the wildtype and *fwd* RNAi embryos at the onset of apical constriction and when apical area of the constriction domain reduces to ∼ 50%. Cyan and yellow boxes mark cells located at the middle of the constriction domain that were typically used for analysis in B – F and G – K, respectively. Scale bars, 20 μm. **(B)** Zoom-in view of cells inside cyan box in (A). Scale bars, 5 μm. **(C-D)** Cell area (C) and anisotropy (D) of individual constricting cells at ∼ 50% apical area reduction. 723 constricting cells from 10 wildtype embryos and 648 constricting cells from 9 *fwd* RNAi embryos were plotted. n.s.: not significant. Two tailed, unpaired Student’s t test. **(E-F)** Standard deviation of cell area (E) and cell anisotropy (F) for constricting cells. Same cells as in C and D were plotted. **: p < 0.01; ***: p < 0.001. Two tailed, unpaired Student’s t test. **(G)** Zoom-in view of cells inside yellow box in (A). Scale bars, 5 μm. **(H-I)** Cell area (H) and anisotropy (I) of individual cells in the middle of the constriction domain at the onset of apical constriction. 242 constricting cells from 7 wildtype embryos and 139 constricting cells from 4 *fwd* RNAi embryos were plotted. n.s.: not significant. Two tailed, unpaired Student’s t test. **(J-K)** Standard deviation of cell area (J) and cell anisotropy (K) for constricting cells. Same cells as in H and I were plotted. n.s.: not significant. Two tailed, unpaired Student’s t test.

Since the reduction of apical cell area during ventral furrow formation is driven by apical Myosin II contractions, we next examined whether knockdown of *fwd* led to any change in apical Myosin II organization during apical constriction. The apical activation and accumulation of Myosin II in *fwd* RNAi embryos were comparable to wildtype control (Supplementary Figure 2, T=0-6 min), but strikingly, we found that Myosin II organized into ring-like structure within the apical domain of each constricting cell, in contrast to the interconnected supracellular network appearance in wildtype embryos (Supplementary Figure 2, T=6-9 min). The mechanistic links between the myosin ring phenotype, the reduced rate of apical constriction, and the loss of heterogeneity in apical cell shape remain to be further elucidated.

### Fwd facilitates apical surface expansion in the non-constricting cells adjacent to the constriction domain

In addition to the constricting cells, depletion of Fwd also affected cell surface expansion in the non-constricting cells outside of the constriction domain. During apical constriction in control embryos, the apical domain of the non-constricting lateral mesodermal cells adjacent to the constriction domain (“flanking cells”) was pulled by the constricting cells and became stretched along the mediolateral direction (Figure 6A, B). In the *fwd* RNAi embryos, however, the flanking cells appeared much less stretched, and this difference, albeit less obvious, can also be seen in the more laterally localized ectodermal cells (Figure 6A, A’, A’’). In line with these observations, quantification of apical cell morphology in the non-constricting cells at 50% apical constriction revealed smaller apical cell area and lower cell anisotropy in *fwd* RNAi embryos compared to the controls (Figure 6B - D). This morphological difference was further confirmed as we compared the average size of the most stretched cells at each medial-lateral row of cells (Figure 6H - J). Of note, the differences in cell area and anisotropy did not exist before the onset of apical constriction (Figure 6E - G). Together, these observations demonstrate that the expansion of the apical domain in the non-constricting cells, which is caused by mechanical stretching from the constriction domain, is impaired in *fwd* RNAi embryos. The distinct phenotype in the constricting and non-constricting cells raised an interesting possibility that Fwd is required for promoting cell surface expansion at both the lateral and apical cell surfaces. However, the mutant phenotype manifests differently in different cell groups, perhaps depending on how cells change their shape and where surface expansion is triggered.

**Figure 6.**
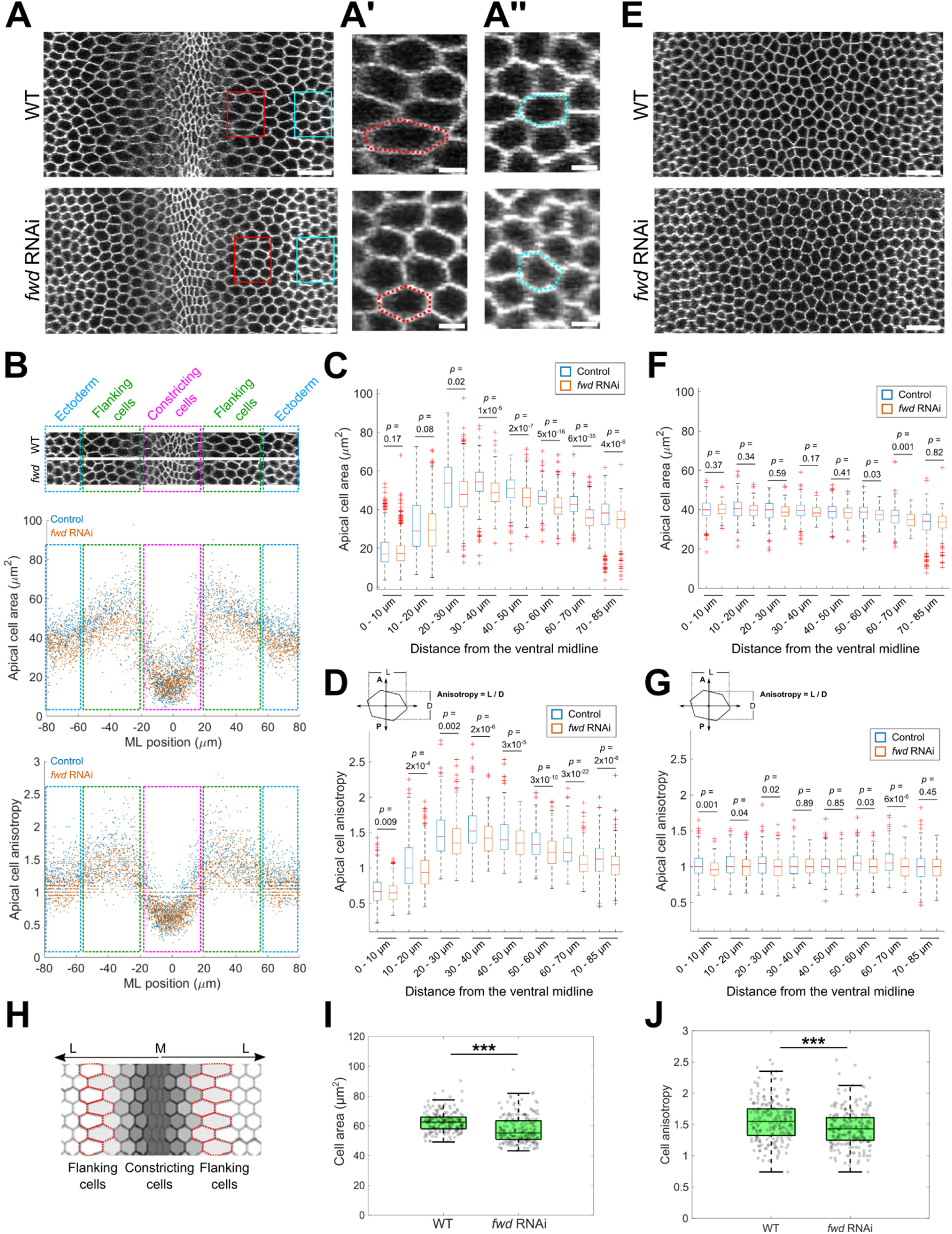
Flanking cells are less stretched in *fwd* RNAi embryos than in the control embryos when similar degree of apical constriction is achieved. **(A)** Surface view of wildtype and *fwd* RNAi embryos when apical area of the constriction domain reduces to ∼ 50%. Scale bars, 20 μm. (A’) and (A’’) are the zoom-in view of the flanking cell region and the lateral ectoderm region marked by the red box and cyan box in (A), respectively. Red and cyan dotted lines show examples of one cell in each genotype. Scale bars, 5 μm. **(B)** Cell area (top) and cell anisotropy (bottom) distribution along mediolateral direction at 50% apical constriction. 0 μm marks the ventral midline. Different regions are indicated with colored boxes. Cells from 9 wildtype and 9 *fwd* RNAi embryos were pooled together. **(C, D)** Comparison of cell area (C) and anisotropy (D) between wildtype and *fwd* RNAi embryos at 50% apical constriction for cells located at different distance from the ventral midline. Two tailed, unpaired Student’s t test. **(E)** Surface view of wildtype and *fwd* RNAi embryos before apical constriction. Scale bars, 20 μm. **(F, G)** Comparison of cell area (F) and anisotropy (G) between wildtype and *fwd* RNAi embryos before apical constriction. Cells from 7 wildtype and 4 *fwd* RNAi embryos were pooled together. Two tailed, unpaired Student’s t test. **(H-J)** Apical area and anisotropy of the most stretched cells along the mediolateral at 50% apical constriction. (H) The most stretched cells in each row of cells (red outlines) that are quantified in (I) and (J). (I) Apical cell area. (J) Apical cell anisotropy. Wildtype: N=203 cells from 9 embryos; *fwd* RNAi: N=210 cells from 9 embryos. ***: p < 0.001. Two tailed, unpaired Student’s t test.

### Fwd knockdown results in slower invagination and abnormal furrow morphology

Given the important role of Fwd in regulating individual cell shape during apical constriction, we examined how Fwd knockdown would affect ventral furrow formation at the tissue scale. We found that depletion of Fwd affected furrow invagination in various aspects (Figure 7A). First, we observed a consistent difference in invagination kinetics between control and *fwd* RNAi embryos (Figure 7B). In the control embryos, invagination occurred first slowly during the first ten minutes, followed by an acceleration of invagination (Figure 7B). The slow and fast invagination phases corresponded to the lengthening and shortening phases of ventral furrow formation, respectively (Supplementary Figure 3A-B) ^38^. In *fwd* RNAi embryos, the transition between the two invagination phases still correlated with the transition from cell lengthening to shortening, but the lengthening phase was ∼46% longer compared to the control embryos (Lengthening phase duration in control: 9.4±1.4 minutes, mean±s.d., N=6 embryos; lengthening phase duration in *fwd RNAi*: 15.1±1.5 minutes, mean±s.d., N=4 embryos; Supplementary Figure 3B). In addition, compared to the wild type, *fwd* RNAi embryos displayed lower rate of furrow invagination during the shortening phase and a moderate reduction in the final furrow depth (Figure 7B, C).

**Figure 7.**
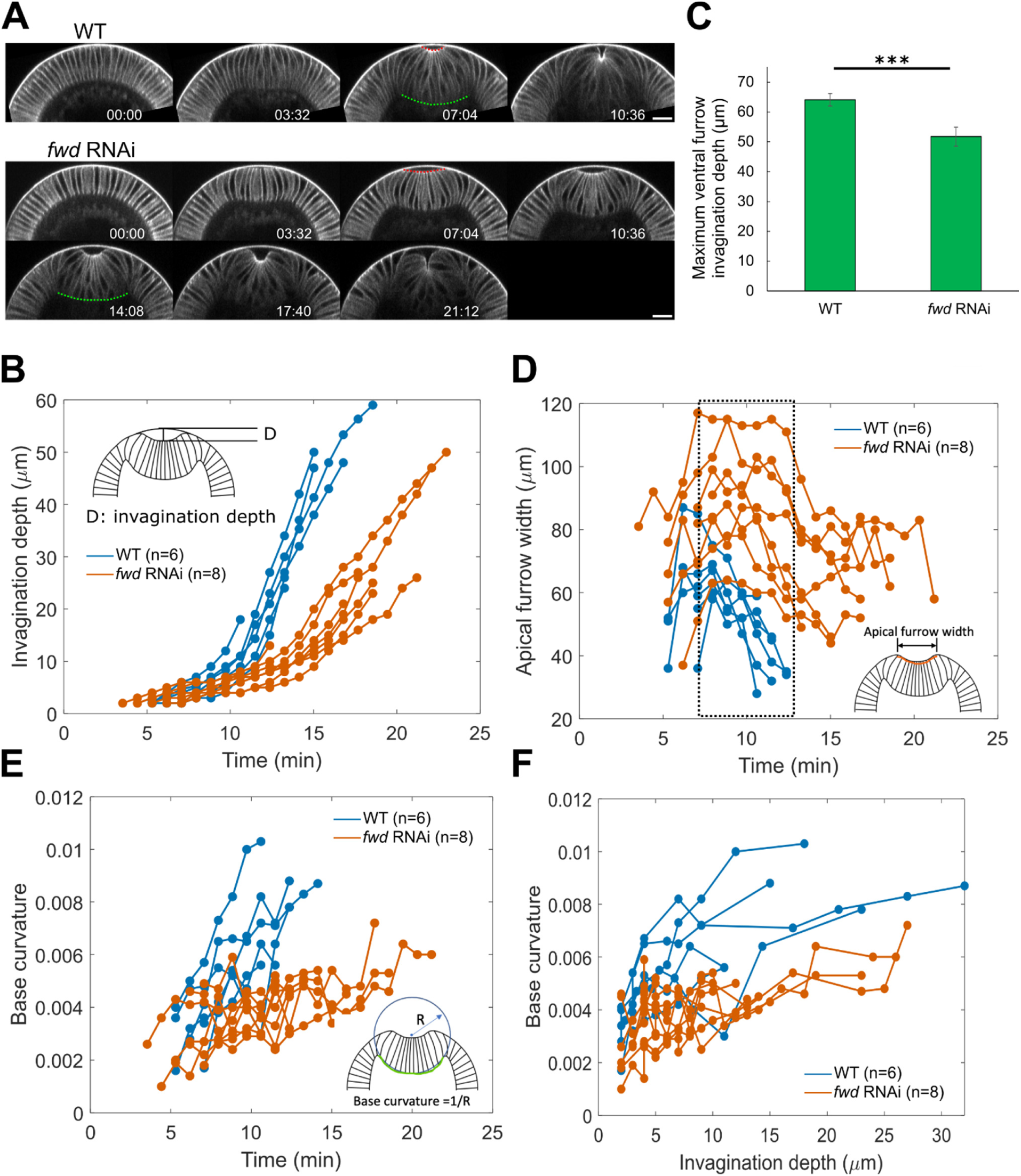
Depletion of Fwd results in slower ventral furrow invagination and abnormal furrow morphology. **(A)** Representative movie stills showing the cross-section view of ventral furrow for wildtype and *fwd* RNAi embryos. Scale bars, 20μm. **(B)** Invagination depth over time. For all the quantifications in this figure, N=6 embryos for wildtype, N=8 embryos for *fwd* RNAi embryos unless otherwise mentioned. **(C)** Maximum invagination depth. N=4 embryos for both wildtype and *fwd* RNAi embryos. Error bars stand for s.d., ***: p < 0.001. Two tailed, unpaired Student’s t test. **(D)** Apical furrow width over time. Dashed box indicates a prolonged phase with wide open furrow in *fwd* RNAi embryos. **(E)** Furrow base curvature over time. **(F)** Furrow base curvature as a function of invagination depth. Furrow base curvature is higher in the control embryos than in *fwd* RNAi embryos at comparable invagination depth.

In addition to the defects in the kinetics of furrow invagination, the furrow morphology was also altered in *fwd* RNAi embryos. The initial phase of ventral furrow formation was comparable between the control and *fwd* RNAi embryos. However, the morphology of the furrow started to deviate between control and *fwd* RNAi embryos as apical constriction progressed (Figure 7A). In both genotypes, a shallow apical indentation was generated approximately 5-7 minutes after the onset of apical constriction and quickly widened up to reach its maximal width. In wildtype embryos, the generation of this wide apical opening was followed by a rapid narrowing of the opening as the furrow folded up and invaginated. In contrast, in the *fwd* RNAi embryos, the apical indentation widened to a larger extent and stayed at the wide configuration for a prolonged time (Figure 7D, dashed box). Of note, the wider apical opening was not due to an expansion of the constriction domain, as the number of constricting cells was comparable between the two genotypes (Supplementary Figure 3C, D). The *fwd* RNAi embryos also showed morphological abnormalities at the basal side of the intermediate furrow near the end of the lengthening phase. Specifically, the basal side of the furrow was flatter in the *fwd* RNAi embryos compared to the control embryos at equivalent invagination stages (Figure 7A, T = 07:04 in the wild type and T = 14:08 in the *fwd* RNAi embryos, Figure 7E, F). Previous studies have shown that the reduction in basal myosin level is important for the cells to expand their base to facilitate furrow invagination (Polyakov et al., 2014; Krueger et al., 2018). We found that the extent of basal myosin loss was comparable between the control and *fwd* RNAi embryos despite the difference in basal curvature, indicating that the flatter base of the intermediate furrow was not caused by defects in basal myosin loss (Supplementary Figure 3E). Taken together, our results demonstrate that depletion of Fwd causes specific tissue-scale abnormalities in ventral furrow formation, including reduced efficiency of furrow invagination and altered intermediate and final furrow morphologies.

### Computer modeling suggests that restricting apical and lateral cell surface expansion may account for different aspects of the ventral furrow phenotype in *fwd* deficient embryos

In order to understand the link between the cell surface expansion defects and the tissue-level abnormalities in *fwd* RNAi embryos, we turned to a modeling approach previously developed by Polyakov et al. ^38^ (Figure 8A; Methods). This 2D vertex model considers the cross-section view of the embryo, where a fully invaginated furrow can be achieved through a combined action of apical constriction, elastic cell cortices (“edges” in 2D) and a non-compressible cell interior. In addition, by adiabatically reducing the basal stiffness (*K_b_*), which mimics the gradual reduction of basal Myosin II during ventral furrow formation, the model can recapitulate different intermediate furrow morphology during the folding process ^38^ (Figure 8B_1_). Finally, the ectoderm in the model undergoes moderate apicobasal thinning, a process that can occur independently of ventral furrow formation in real embryos ^34^. Despite the simplifications on the morphology and mechanical properties of the cells, the model can successfully recapitulate the stepwise changes in tissue morphology during ventral furrow formation and predict the bistable characteristic of the mesoderm ^34, 38^. This modeling framework was advantageous in testing our hypothesis since the outcomes of the model could reveal specific morphological defects at multiple stages of furrow formation. Furthermore, the properties of the apical, basal and lateral cortices could be controlled separately in the model, thereby allowing us to examine the specific impact of restricting apical or lateral surface expansion on furrow formation.

**Figure 8.**
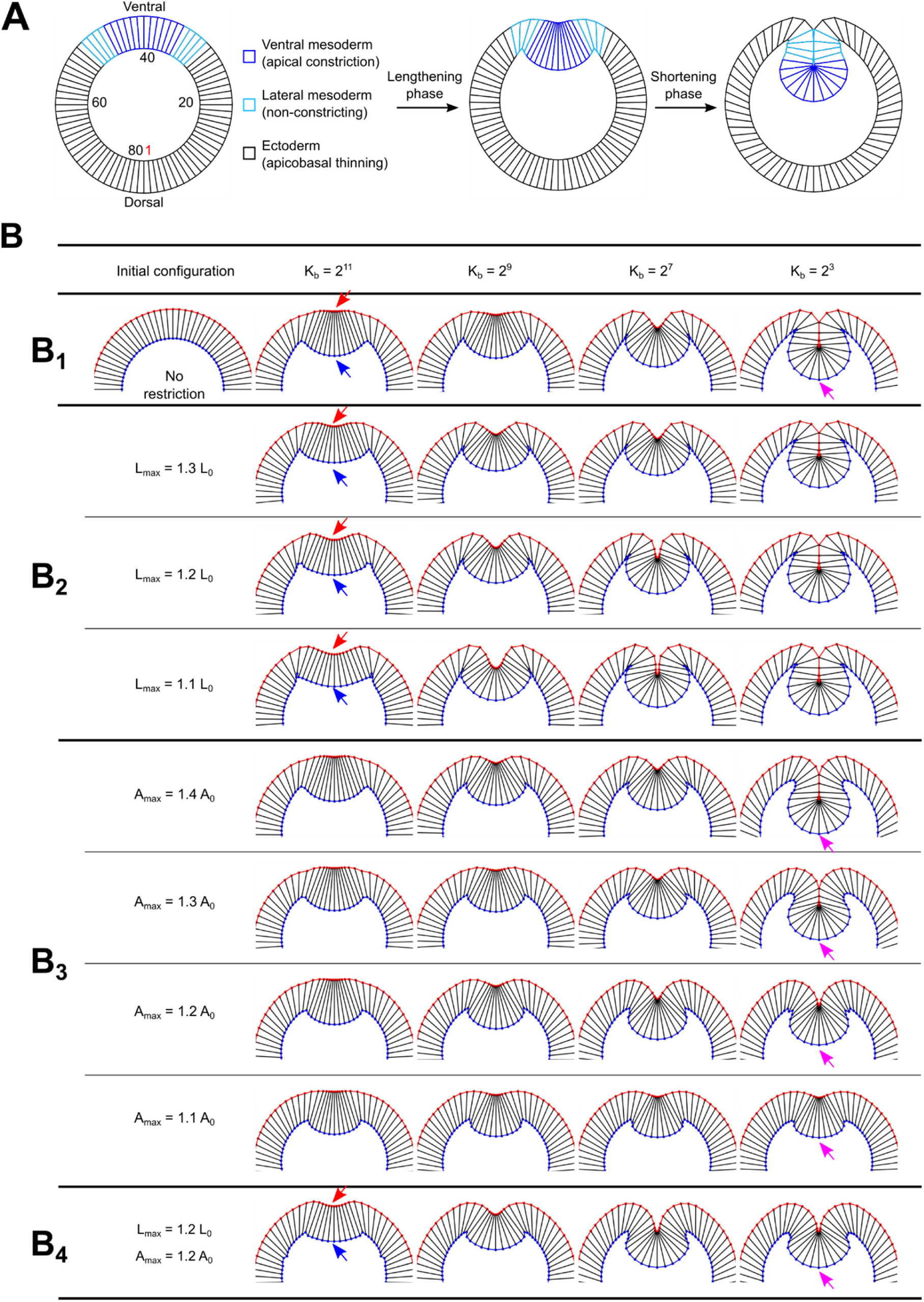
Simulation shows distinct impact of restricting apical and lateral surface expansion on ventral furrow formation. **(A)** 2D vertex model testing the impact of defects in cell surface expansion on ventral furrow formation. The model considers the cross-section view of the embryo, which contains a ring of 80 columnar-shaped cells. In the model, the cortices of cells resist deformations as elastic springs, and the cells have a strong propensity to maintain constant volume. The model is driven out of equilibrium by apical constriction in the ventral mesoderm (the upmost 12 cells). In addition, the ectodermal cells located outside of the mesoderm domain undergo apical-basal shortening of the ectoderm (Methods). **(B)** Simulation results (B_1_ – B_4_). **(B_1_)** When simulating normal ventral furrow formation, the model transitions through a series of equilibrium states governed by a stepwise reduction of basal stiffness (K_b_). **(B_2_)** Ventral furrow formation when the maximal elongation of the lateral membrane (L_max_) is constrained to 1.3-, 1.2- and 1.1-fold of its original length L_0_. As the constraint on L_max_ increases, the morphology of the intermediate furrow becomes increasingly abnormal, featured by a wider apical indentation (red arrows) and a flatter base (blue arrows). On the other hand, the final invagination depth is not affected. **(B_3_)** Ventral furrow formation when the maximal elongation of the apical membrane (A_max_) is constrained to 1.4-, 1.3-, 1.2- and 1.1-fold of its original length A_0_. As the constraint on A_max_ increases, the final furrow depth becomes increasingly smaller (magenta arrows). The intermediate furrow morphology is not substantially affected. **(B_4_)** When both L_max_ and A_max_ are constrained to 1.2-fold of their original length, the effects are additive, and the model shows defects in both intermediate furrow morphology and the final invagination depth, which recapitulate the ventral phenotype in *fwd* deficient embryos.

First, we tested the impact of restricting lateral membrane expansion on ventral furrow formation. To this end, we imposed a constraint on the maximal length of the lateral edges (*l ≤ l_max_,* Methods). Under wildtype conditions, only the cells that undergo apical constriction elongate in the apical-basal direction. Therefore, only these cells would be directly impacted by the constraints on lateral expansion (Supplementary Figure 4A, magenta box). We tested conditions where *l_max_* is 1.3, 1.2 or 1.1 times of the original lateral length *l_0_*. In all three cases, the impose of the constraint on lateral expansion led to a wider furrow opening and a flatter furrow base prior to the lengthening-shortening transition (Figure 8B_2_, red and blue arrows, respectively), which resembles the phenotype observed in *fwd* RNAi embryos. The severity of the phenotype increases as *l_max_* decreases. Despite this defect, the model could still invaginate, and the final depth of the furrow was not affected (Figure 8B_2_).

Using a similar approach, we next examined the impact of restricting apical membrane expansion on ventral furrow formation. In this case, we imposed a constraint on the maximal length of the apical edges (*a ≤ a_max_*; Methods). Under wildtype conditions, the flanking cells in the model showed most prominent apical expansion, up to a factor of 2 (Supplementary Figure 4B, magenta box). In addition, the more laterally localized ectodermal cells also moderately expanded their apical domain, by a factor of ∼ 1.3 (Supplementary Figure 4B, green box). The expansion of the apical surface in these cells offset the loss of apical size in the constriction domain and resulted in a ∼ 15% net increase in the total apical surface size when the furrow was fully invaginated (Supplementary Figure 4C). Interestingly, the model predicted that the impact of restricting apical expansion was distinct from restricting lateral expansion. Instead of affecting the intermediate furrow morphology, restricting apical expansion mainly impacted the final invagination depth. Under conditions where *a_max_* > *1.3a_0_*, when the constraint mostly affected the flanking cells but not the ectodermal cells, furrow invagination appeared normal (Figure 8B_3_, *a_max_* = *1.4a_0_*; Supplementary Figure 4B, C). However, under conditions where *a_max_* < *1.3a_0_*, when both the flanking cells and the ectodermal cells were affected, the final invagination depth was reduced (Figure 8B_3_; Supplementary Figure 4B, C). Of note, an *a_max_* of *1.25a_0_* resulted in a reduction of final invagination depth from 60 μm to 50 μm, which is the level of defect observed in *fwd* RNAi embryos (Supplementary Figure 4C, green arrow).

Finally, we tested the effect of combining the restrictions on apical and lateral expansion. In this case, the simulated ventral furrow showed a combined phenotype, i.e., a wider furrow opening and flatter base at the lengthening-shortening transition and a reduced furrow depth at the end of invagination (Figure 8B_4_). These phenotypes qualitatively resembled the morphological defects of ventral furrow in *fwd* RNAi embryos, suggesting that the tissue-level abnormality in the mutant embryos can be attributed to defects in cell’s ability to expand its surfaces. Our model also predicts that restrictions on apical and lateral surface expansion may have distinct impact on tissue-scale mechanics, which would be interesting to test in the future.

### Fwd and PI4KIIα have partially redundant function in membrane growth during cellularization

Given the important function of exocytic trafficking in plasma membrane expansion, we wondered whether Fwd also regulates other morphogenetic processes in early *Drosophila* embryos that require cell surface expansion. As mentioned above, depletion of Fwd resulted in a moderate reduction in apical-basal cell length at the beginning of gastrulation, which is indicative of cellularization defect. To further analyze this phenotype, we measured cell length over time during cellularization. Previous studies have shown that the ingression of cleavage furrows proceeds in separate slow and fast phases during cellularization ^40, 41^. We found that depletion of Fwd did not significantly affect the rate of furrow ingression during the slow phase, but the rate of furrow ingression during the fast phase was mildly reduced compared to the control embryos (Figure 9A, B). Note that this defect was only observed upon relatively strong depletion of Fwd (Figure 2F-H). The mild cellularization phenotype of *fwd* RNAi embryos prompted us to ask whether other PI 4-kinases share redundant function with Fwd during cellularization. Previous study has shown that another PI 4-kinase, PI4KIIα, also displays Golgi localization in *Drosophila* ^20^. We found that knockdown of PI4KIIα did not result in noticeable defects in cellularization (Figure 9C, D). However, when we knocked down both Fwd and PI4KIIα, the resulting embryos became severely disrupted with no sign of cellularization (Figure 9C). Reducing the expression level of shRNAs in the double knockdown condition prevented the drastic cellularization failure but resulted in substantially reduced cell length at the end of cellularization (Figure 9C, D). This dramatic additive effect suggests that Fwd and PI4KIIα function in a largely redundant manner during cellularization.

**Figure 9.**
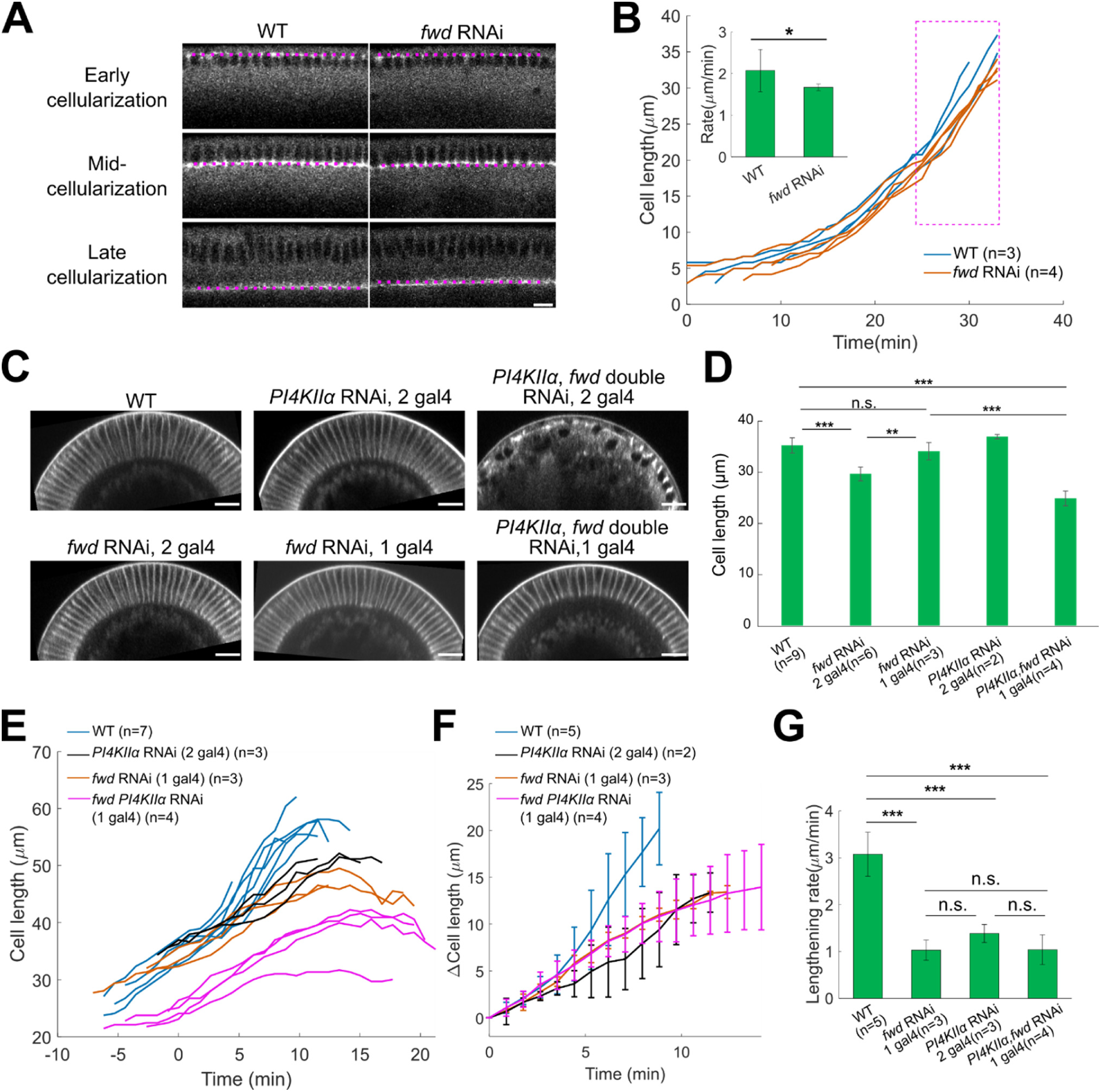
Fwd and PI4KIIα function redundantly in cell membrane growth during cellularization. **(A)** Membrane growth during cellularization in wildtype and *fwd RNAi* embryos. Midsagittal plane images at the dorsal side of the embryo were taken over time. Cellularization front (magenta dotted lines) was visualized by Sqh-mCherry. Scale bar, 10 μm. **(B)** Cell length over time during cellularization. Time 0 is the time point when the cellularization front reaches the top of the nuclear. For embryos missing time 0, embryos are aligned based on the general trend of cleavage furrow ingression. Inset shows the rate of furrow ingression during late cellularization (magenta box). Error bar stands for s.d.; *: p < 0.05; One-sided Wilcoxon rank sum test. **(C)** Representative cross-section view images of embryos from different genetic background at the end of cellularization. Scale bars, 20 μm. **(D)** Quantification of cell length at the end of cellularization for different genetic background. Error bar stands for s.d.; n.s.: not significant; ***: p < 0.001; **: p < 0.01; Two tailed, unpaired Student’s t test. **(E)** The length of the apically constricting cells over time during apical constriction for embryos from the indicated genetic backgrounds. 0 min represents the onset of gastrulation. **(F)** Average length of the apically constricting cells over time during apical constriction. Error bar stands for s.d.. **(G)** Rate of cell lengthening at phase 2. Error bar stands for s.d.. n.s.: not significant; ***: p < 0.001; Two tailed, unpaired Student’s t test.

Next, we asked whether functional redundancy between Fwd and PI4KIIα also exists during cell lengthening in ventral furrow formation. Although PI4KIIα single knockdown did not obviously affect cellularization, when these mutant embryos entered gastrulation, they showed similar defects in cell lengthening as in the *fwd* deficient embryos (Figure 9E-G). Interestingly, when the double knockdown embryos entered gastrulation, despite the reduced starting cell length, the rate and extent of cell lengthening were comparable to those in the *fwd* single knockdown embryos (Figure 9E-G). Taken together, our results revealed important function of Fwd and PI4KIIα in cell surface expansion during cellularization and apical constriction-mediated cell lengthening. Interestingly, while the functions of the two enzymes are largely overlapped during cellularization, their roles in cell lengthening appear to be less redundant.

## Discussion

In this study, we identified *Drosophila* PI4KIIIβ homologue, Fwd, as an important regulator of epithelial morphogenesis in early *Drosophila* embryos. During ventral furrow formation, disruption of Fwd function resulted in a reduction in the rate of cell lengthening and cell surface area increase in cells undergoing apical constriction. Depletion of Fwd also resulted in alterations in the spatial organization of apical myosin and a moderate reduction in the rate of apical constriction, but this phenotype is separable from the defects in cell lengthening and surface expansion. In addition to the constricting cells, the non-constricting cells adjacent to the constriction domain also displayed impaired surface expansion, as revealed by a reduced stretching of their apical domain compared to the wildtype embryos. Finally, during cellularization, simultaneous disruption of Fwd and another PI 4-kinase, PI4KIIα, severely affected cell surface expansion mediated by the ingression of cleavage furrows. Together, these results provide, to the best of our knowledge, the first description of the role of PI 4-kinases in promoting cell surface expansion and cell shape change during epithelial morphogenesis. Using computer modeling, we further demonstrated that restricting apical and lateral surface expansion can lead to specific defects in ventral furrow morphology that closely resemble the phenotypes observed in *fwd* deficient embryos. These findings point to a potential mechanistic link between cell surface “expandability” and tissue-scale mechanics during epithelial folding.

Previous studies have shown that Fwd plays an important role in meiotic cytokinesis during spermatogenesis ^31, 32, 42^. In *fwd* mutant males, contractile rings can still form and constrict in dividing spermatocytes, but cleavage furrows are unstable and later retract. Fwd is known to regulate the Golgi to plasma membrane trafficking, and similar cytokinesis defects have also been observed in mutants of other Golgi associated proteins ^31, 42–44^. These observations suggest that the spermatocyte cytokinesis defects in *fwd* mutant are caused by defects in membrane trafficking. In this work, we showed that Fwd also contributes to membrane growth during cellularization, an atypical cytokinesis process, although in this case Fwd functions in a largely redundant manner with PI4KIIα. Given the observation that *fwd* mutants do not show obvious defects in regular mitotic cytokinesis ^32^, an interesting future investigation is to determine the relative contribution of Golgi-localized PI 4-kinases in different types of cytokinesis processes.

While numerous studies have shown the important role of exocytosis in membrane growth during cytokinesis, including cellularization ^45, 46^, the role of exocytic trafficking in cell membrane expansion in support of rapid cell shape change during epithelial remodeling is less well understood. We show that during ventral furrow formation, apical constriction-mediated cell lengthening involves a prompt increase in the cell surface area. Our identification of Fwd, a PI4 kinase important for Golgi-PM trafficking, as a regulator of cell lengthening further suggests that the observed cell surface expansion involves active exocytic membrane insertion. Future experiments directly monitoring new membrane addition during ventral furrow formation would be important to further test this hypothesis.

So far, the only reported *fwd* phenotypes associated with surface area regulation are spermatocyte cytokinesis defects initially discovered by Brill et al ^32^ and the cellularization and gastrulation defects described in this study. A common characteristic shared between these processes is that all of them involve rapid membrane expansion, which likely imposes a high demand on intracellular membrane supply. In spermatocytes, meiotic divisions occur in rapid succession, causing about 60% increase in surface area in less than two hours ^31^. Likewise, during cellularization, the cells increase their surface area 25 fold in about one hour ^40, 47^. During apical constriction-mediated cell lengthening, the cell surface area increases by ∼25% in about eight minutes. These observations raise the hypothesis that the function of Fwd becomes more important when there is an acute demand for membrane supply. An interesting future question is whether the activity of Fwd is subjected to regulation by the elevated demand on intracellular membrane supply and, if so, how does the cell sense such demand. Another important future question is how Fwd executes its function to facilitate cell surface expansion during cellularization and cell lengthening. Two modes of PI4KIIIβ function have been discovered in animal cells, both of which can promote exocytic membrane insertion. PI4KIIIβ can serve as a docking site to recruit Rab11 to TGN through direct protein-protein interaction, which has been observed both in vitro and in vivo ^28, 30^. Alternatively, PI4KIIIβ catalyzes the production of PI(4)P lipids at TGN. PI(4)P helps to recruit various proteins important for trafficking and can also indirectly impact trafficking through affecting other pools of membrane lipid species ^22–26^. Understanding the mode of action will be a key step to uncover the relevant downstream molecular network regulating cell surface area during epithelial remodeling.

Our work also shed light on the impact of restricting cell surface expansion on tissue-level remodeling during morphogenesis. During ventral furrow formation, disrupting the function of Fwd resulted in prolonged lengthening phase, aberrant intermediate furrow morphology prior to invagination, slower furrow invagination and shallower furrow at the end of invagination. While a complete understanding of the cause of these phenotypes and their potential connections awaits further investigation, our modeling analysis suggests that certain key aspects of the tissue-level morphological abnormality in the mutant embryos can be attributed to defects in cell surface expansion. It is important to note that in *fwd* deficient embryos, cell surface expansion is attenuated, but not completely inhibited, which may explain the relatively moderate phenotype on ventral furrow formation. Future investigations with novel approaches that allow acute and more complete block of cell membrane expansion will offer a further test on this point. In addition, new modeling approaches that contain the temporal components, which is lacking in our current modeling framework, will help us to start to understand how restricting cell surface expansion might contribute to the defects in tissue folding kinetics observed in *fwd* deficient embryos.

## Materials and methods

### Fly stocks and genetics

*Drosophila melanogaster* flies were grown and maintained at 18°C and crosses were maintained at room temperature (21 – 23 °C). All flies were raised on standard fly food. For embryo collection, flies with corresponding genotype were used to set up cages and maintained at 18°C, and embryos were collected from apple juice agar plate containing fresh yeast paste.

The UAS-shRNA lines targeting *fwd* (TRiP *fwd,* BDSC stock#35257) and *PI4KIIα* (TRiP *PI4KIIα*, BDSC stock#65110) were obtained from Bloomington Drosophila Stock Center. A TRiP *PI4KII*; TRiP *fwd* stock were generated from cross for the double RNAi experiments. The *fwd^3^* mutant stock (*fwd^3^*/TM6) was a gift from Julie Brill and have been described previously (Brill et al., 2000; Polevoy et al.,2009). *fwd*^3^ /TM6 flies were crossed to flies from a deficiency line covering the *fwd* gene locus (Df(3L)Exel9057/TM6B, BDSC stock#7920) to generate loss of function *fwd* mutant.

The candidate RNAi and dominant negative lines used for the screen for lengthening defects and the phenotypes in ventral furrow formation was listed in Table 1. The genes included exocytic trafficking related small GTPases and their regulators and/or effectors (Rab4, Rab8, Rab11, Arf1, Sec71, Garz, Nuf, Crag, Brun, Gga), motor proteins (dynein, Myosin V), lipid regulators (Cert, PI4KIIα, PI4KIIIα, Fwd, Sac1) and components of exocyst complex (Sec5). For some small GTPases, the impact of expression of dominant negative form of the protein was also examined ^48, 49^.

For RNAi-mediated knockdown, female flies from RNAi lines were crossed to male flies from GAL4 driver lines, and embryos from F1 flies were collected for imaging. GAL4 driver lines carrying matα4-GAL-VP16 (denoted as “mat67” on the 2nd chromosome and “mat15” on the 3rd chromosome) were used to drive maternal expression of shRNA in the embryo ^50^. For control experiment, wildtype female Oregon R flies were crossed to the GAL4 driver lines. For examining the cellularization and lengthening defects, the driver line stock mat67 Sqh::mCherry; mat15 Ecad::GFP was used for knockdown experiments with 2 copies of GAL4, and the driver line stock Sp/Cyo; mat15 Ecad::GFP was used for knockdown experiments with single copy of GAL4. Unless otherwise specified, all the knockdown experiments are performed with 2 copies of GAL4.

To examine myosin in *fwd* RNAi background, female flies from Oregon R (control) and TRiP *fwd* stock were crossed to male flies containing Sqh::mCherry and 2 copies of maternal GAL4 drivers. Embryos from F1 flies were collected for imaging.

### Live imaging

Embryos were dechorionated in 40% bleach (∼3% Sodium Hydrochloride), rinsed with water 12 times and mounted in water with the ventral side facing up in a 35 mm MatTek glass-bottom dish (MatTek Corporation). All live imaging was conducted on an upright Olympus FV-MPERS multiphoton microscope equipped with the InSight Deepsee Laser System, an Olympus 25×/1.05 water dipping objective (XLPLN25×WMP2) and Fluoview software. 920 nm laser was used to excite GFP/YFP. Unless otherwise mentioned, a 512 × 512 pixel (pixel size: 0.331 μm/pixel) of region of interest was imaged using resonant scanner with frame average of 16 times. Embryos were imaged from the ventral surface to 80 μm below surface with a Z step size of 1 μm and temporal resolution of 53 sec/frame. For movies used for 3D reconstruction, in order to gain higher signal-to-noise ratio, a 300×150pixel (pixel size: 0.331 μm/pixel) of region of interest were imaged using galvanometer scanner with frame average of 4 times. The depth and step size in Z was the same while the temporal resolution was set as 2 min/frame to minimize photobleaching.

### Image processing and analysis

All images were processed using ImageJ (NIH) and MATLAB (MathWorks). Embryos were aligned based on the onset of gastrulation when ventral mesodermal precursor cells start to constrict apically. E-cadherin-GFP was imaged as a membrane marker for all image analyses described below.

#### 2D analysis of cell lengthening (cross-section view)

For analysis of the rate of cell lengthening during apical constriction, an average projection of 20 slices of cross-section images was first generated, and the apical-basal height of the cell located at the ventral midline was measured in ImageJ. For analysis of the rate of apical constriction, a row of ∼10 cells in mediolateral direction centered around ventral midline was selected from the en face view images and tracked over time. For each time point, the farthest left and right boundary points at the most apical side of this row of cells were determined and mapped onto the cross-section view images. Then, the length of the apical curve between the left and right boundary points was measured from the cross-section view using segmented line tool in ImageJ. To calculate the constriction rate and lengthening rate, the apical domain width over time plot and cell length over time plot were first smoothed and interpolated (from 2-min interval to 1-min interval), and the derivative of the resulting curves were computed to determine the rate of change.

#### 3D analysis of cell lengthening

3D segmentation and quantification of individual cells were performed as previously described ^34^. Individual cells near the ventral midline were manually tracked over the course of ventral furrow formation and segmented using the Carving tool of the “Ilastik” program ^51^. For each timepoint, a Z-stack of images covering the entire depth of the ventral tissue was used as the input. Manual corrections were performed for the apical and basal most regions of the cells, where the automatic segmentation by Ilastik was usually not optimal. Manual corrections were carried out by manually outlining the cell at the relevant Z-planes using the multi-point took in ImageJ. The resulting measurements were then incorporated into the automatic segmentation using a custom MATLAB script to create the final 3D rendering of the cell. Three control embryos and three *fwd* RNAi embryos were analyzed. For each embryo, 3 – 6 cells were segmented over the course of the lengthening phase at 2-minute intervals.

A custom MATLAB script was used to process the reconstructed cells and analyze the cells’ surface area, volume, cell length along the apical-basal axis, and apical area. Surface area and volume were measured using the “surfaceArea” and the “volume” functions in MATLAB, respectively. To measure the apical-basal cell length and apical area, the 3D cell mask was rotated so that the cell apical-basal axis became vertical. This was followed by regenerating the Z stack of 2D slices. The cell length was calculated by summing up the distances between the centroids of cell slices at neighboring Z-planes except for the apical-most (3 μm) and basal-most (3 μm) regions, where the distances were determined by directly measuring the vertical height to avoid measurement errors due to irregular shapes in these regions. The apical area was determined by measuring the average area of the apical slices, ranging from 1 – 2 μm from the apical surface of the rotated cell mask.

#### Analysis of apical shape and area change (surface view)

For quantification of the apical cell shape, the image stack was first corrected for any tilting in the mediolateral axis using ImageJ such that the ventral midline was positioned near the center of the image. Next, a custom MATLAB script was used to generate a flattened surface view of the embryo that accounted for the curvature of the embryo. Next, individual cell was segmented from the surface view (2D) using Cellpose, a deep learning-based segmentation software ^37^. A custom MATLAB script was then used to extract and calculate relevant parameters such as cell position, apical area and apical anisotropy. To identify the most-stretched cells from the surface view, a set of evenly spaced mediolateral sampling lines were generated, and among cells falling on each sampling line, cells with maximal area on left and right side of the ventral midline were identified as the most stretched cells.

### Energy minimization-based 2D vertex model for ventral furrow formation

The energy minimization-based 2D vertex model for ventral furrow formation was constructed as previously described ^34, 38^. The model considers the cross-section view of the embryo, which contains a ring of 80 columnar-shaped cells that resembles the number and geometry of the primary embryonic epithelium formed during cellularization. The morphology of the cells is maintained by the following mechanisms. First, the apical, basal and lateral membranes of the cells resist stretching or compressing like elastic springs. Second, the cells resist changes in cell volume (area in the 2D model). The model is driven out of the initial energy equilibrium by exerting the following two “active” forces. First, the apex of the cells in the ventral region of the model embryo has a propensity to shrink, which resembles apical constriction. Second, the apical-basal length of the ectodermal cells in the model embryo has a propensity to reduce, which resembles ectodermal shortening process observed in real embryos during gastrulation. Upon application of these active forces, the model transitions through a series of intermediate equilibrium states by adiabatically reducing basal spring stiffness, which recapitulates basal myosin loss during ventral furrow formation. These intermediate equilibrium states resemble the intermediate furrow morphology observed during the lengthening-shortening transition in the real embryo.

The energy equation that describes the mechanical properties of the model is given by the following expression:

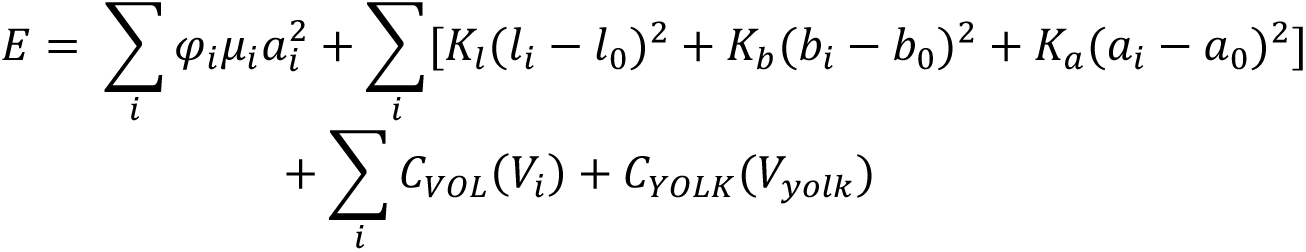

The first term in the equation describes the active apical forces that mediate apical constriction. The term *φ_i_µ_i_* sets up the spatial distribution of apical contractility. *φ_i_* equals to 1 for mesoderm cells (18 cells, with 9 cells flanking each side of the ventral midline) and 0 for the cells outside of the mesoderm domain (“ectodermal cells”). *µ_i_* is a Gaussian function that centers at the ventral midline. Specifically,

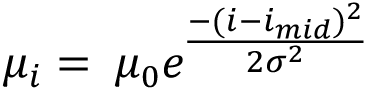

Here *i_mid_* is the cell ID at the ventral midline, *µ_0_* sets up the strength of apical constriction and *σ* sets up the width of the Gaussian function. Of note, the Gaussian function of apical force distribution in the mesoderm domain is set in such a way that only 12 cells at the ventral most region of the embryo will undergo apical constriction, mimicking the situation in real embryos.

The second term in the equation describes the elastic resistance of the membrane springs. *a_i_*, *b_i_*, and *l_i_* are the length of the apical, basal and lateral springs of cell *i*, respectively. The resting length of these springs, *a_0_*, *b_0_*, and *l_0_*, respectively, are set to be equal to the initial length of the springs. The spring constant of the apical, basal and lateral springs are given by *k_a_*, *k_b_* and *k_l_*, respectively. Note that in order to drive apical-basal shortening of the ectoderm in the model, the resting length of the lateral springs (*l_0_*) of the cells outside of the mesoderm domain is set to be 80% of their original length, as described previously ^34^.

The last two terms, *C_VOL_* and *C_YOLK_*, describe the constraint due to volume (“area” in 2D) conservation of the cells and the yolk, respectively. When the volume deviates from the resting value, penalties are imposed as described by the following equations:

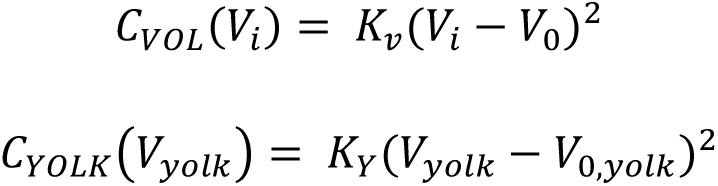

Here, *V_i_* is the volume of cell *i*. *V_0_* is the initial equilibrium volume of cell *i*. Similarly, *V_yolk_* is the volume of the yolk. *V_0,yolk_* is the initial equilibrium volume of the yolk. *K_v_* and *K_yolk_* set the degree of the penalty when the cell volume and the yolk volume deviate from the initial equilibrium values, respectively.

The following approach is used to implement the constraint on lateral membrane expansion. When the lateral spring reaches a certain threshold *l_max_*, a strong penalty is implemented for any further stretching of the spring. The penalty is defined as:

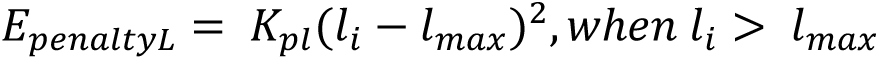

In the simulation, the following thresholds have been tested: *1.3l_0_*, *1.2l_0_*, and *1.1l_0_*. *K_pl_* is set as a constant of 100,000. Note that although all cells in the model are subjected to this constraint, the constraint only affected the apically constricting cells due to the way how cells change their apicalbasal length during ventral furrow formation.

The constraint on apical membrane expansion is implemented in a similar manner. The penalty for exceeding the threshold apical spring size, *a_max_*, is given by:

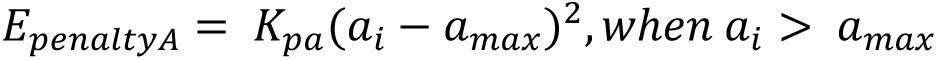

The thresholds for apical spring length tested in the simulation are *1.3a_0_*, *1.2a_0_*, and *1.1a_0_*. *K_pa_* is set as a constant of 100,000.

List of parameters used in the simulation:

| *Parameter* | *Value* |
| --- | --- |
| $K_a$ | 30 |
| $K_l$ | 20 |
| $K_b^*$ | $2^{11} \rightarrow 2^0$ |
| $\mu_0$ | 2000 |
| $\sigma$ | 3 |
| $K_v$ | 5000 |
| $K_Y$ | 1 |

\*: In the simulation, $K_b$ decreases adiabatically from $2^{11}$ to $2^0$ , with a step size of 2-fold reduction.
Accordingly, the model transition through a series of energy-minimum equilibrium state defined by each value of $K_b$ . These equilibrium states represent the intermediate and final furrow morphology.

### Statistics

Statistical comparisons were performed using two-tailed Student’s t tests after Shapiro-Wilk normality test. Sample sizes can be found in the figure legends.

## Data availability

The original data generated in this work are available upon request.

## Code availability

All computer codes used in this study are available upon request.

## Acknowledgments

We thank members of the He lab and the Griffin lab for discussions throughout this work; Yashi Ahmed and Charles K.Barlowe for valuable suggestions on this project; Ann Lavanway and Zednek Svindrych for technical support with imaging. We thank the Brill lab for sharing reagents. We thank the Bloomington *Drosophila* Stock Center (NIH P40OD018537) for fly stocks. This study was supported by NIGMS ESI-MIRA R35GM128745 and American Cancer Society Institutional Research Grant #IRG –82-003-33 to B.H. The study used core services supported by STANTO15R0 (CFF RDP), P30-DK117469 (NIDDK P30/DartCF), and P20-GM113132 (bioMT COBRE).

## Author Contributions

W.C. and B.H. designed the study. W.C. performed the experiments. W.C., V.B and B.H. analyzed the data. W.C. wrote the first draft of the manuscript. All authors contributed to the final version of the manuscript.

