## Supplementary Figures for "PI 4-kinases promote cell surface expansion and facilitate tissue morphogenesis during *Drosophila* cellularization and gastrulation"

#### Supplementary Materials

##### Supplementary Figure 1 - 4

### Supplementary Figure 1

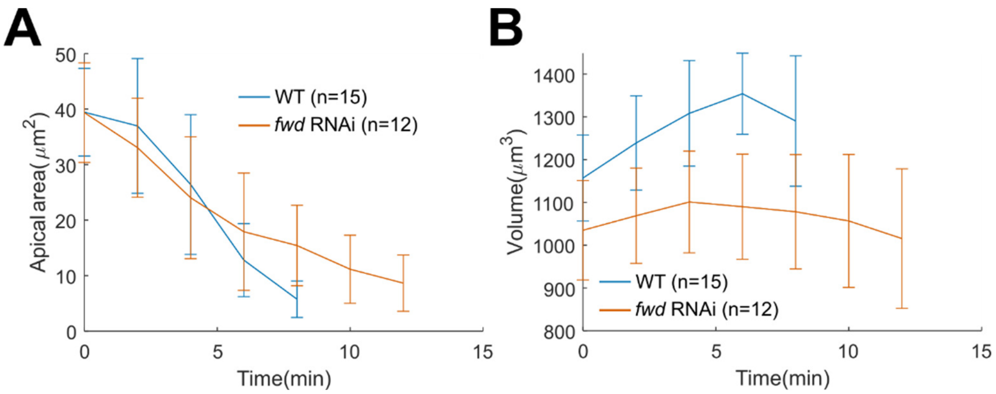

**Supplementary Figure 1. Measurement of apical area and cell volume in 3D segmented cells.**

**(A, B)** Average apical area (A) and cell volume (B) over time in wildtype and *fwd* RNAi embryos during the lengthening phase of ventral furrow formation. Error bars stand for s.d. Same data as in Figure 4 are used.

**Supplementary Figure 2**

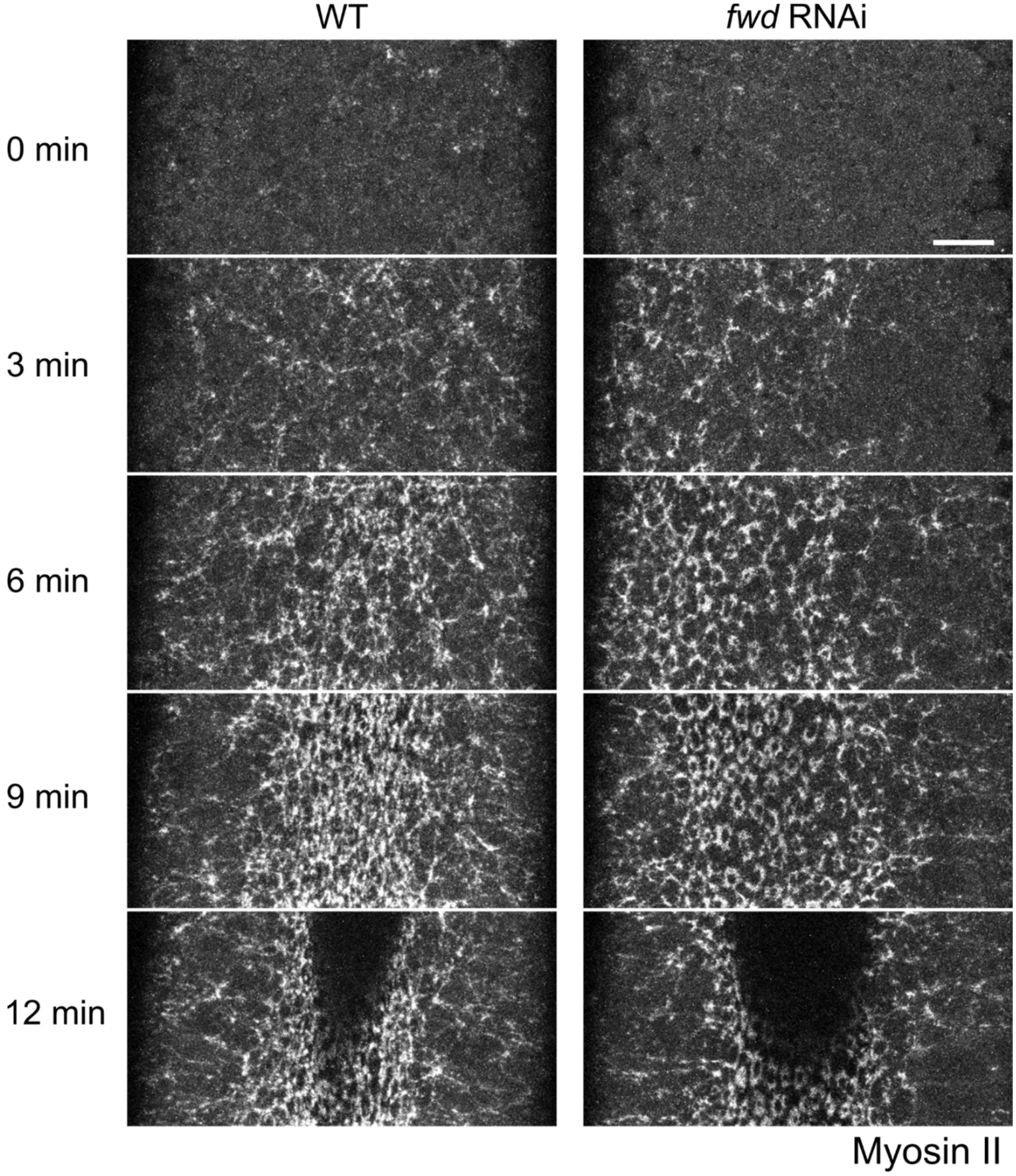

**Supplementary Figure 2. Depletion of Fwd results in apical myosin ring formation**

In *fwd* RNAi embryos (N=5 embryos), apical Myosin II forms individual ring-like structure in each cell apex instead of forming a supracellular network. Scale bar, 10 μm.

### Supplementary Figure 3

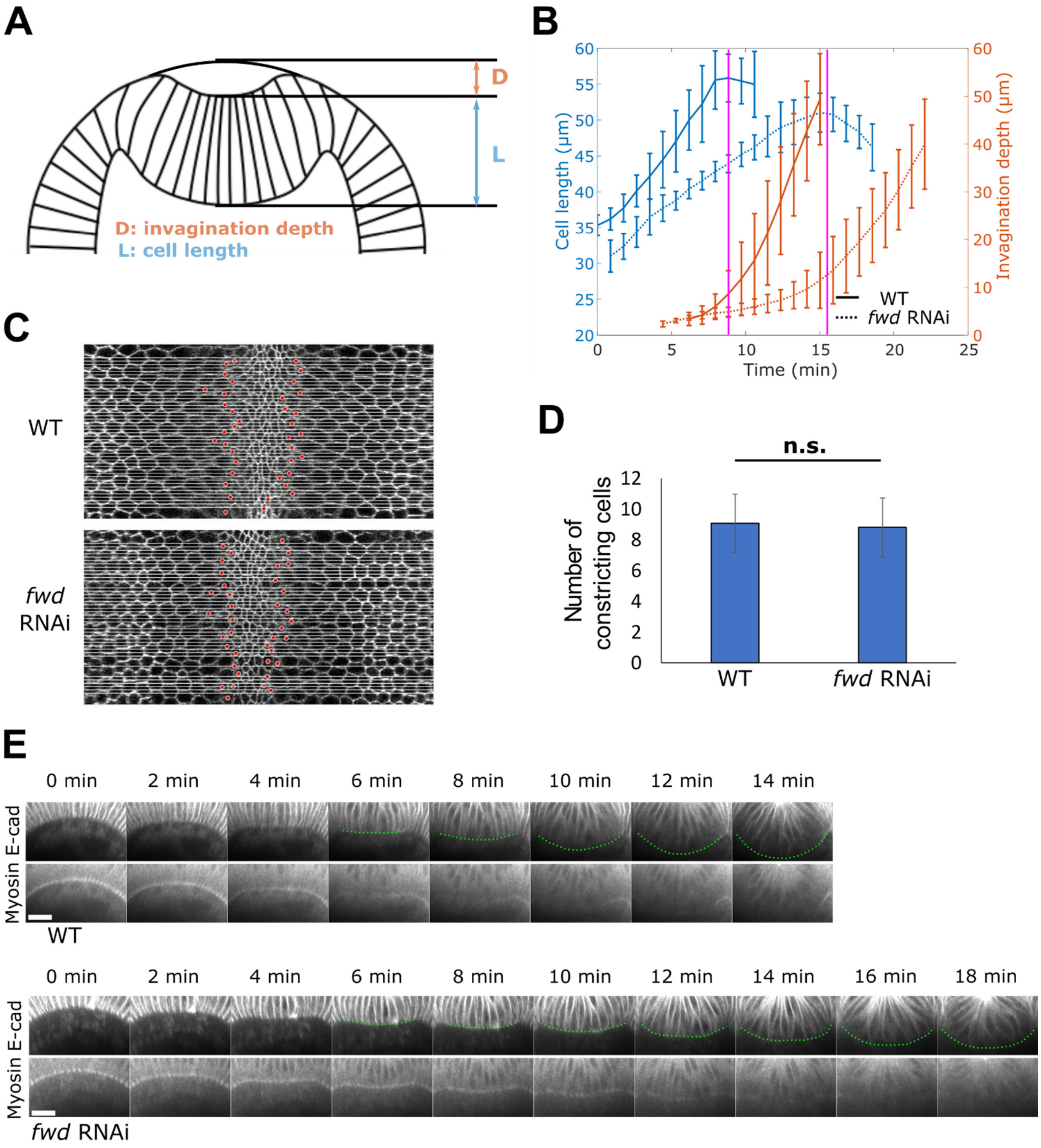

**Supplementary Figure 3. Depletion of Fwd leads to ventral furrow formation defects despite the normal size of the constriction domain and normal loss of basal myosin over time.**

**(A)** Cartoon diagram showing measurements for invagination depth and cell length.

**(B)** Cell length and invagination depth over time. In both wildtype and *fwd* RNAi embryos, the transition point from lengthening to shortening phase corresponds to the turning point from slow invagination and fast invagination phase (magenta lines). Error bar stands for s.d.. N=6 embryos for wildtype, N=4 embryos for *fwd* RNAi embryos.

**(C-D)** Quantification of the number of constricting cells. (C) Surface view of embryos when apical area reduction is ~50%. For both wildtype and *fwd* RNAi embryos, a time point with similar apical constriction was selected. Cells with area smaller than its beginning area at onset of apical constriction were counted as constricting cells. Evenly spaced mediolateral horizontal lines (white lines) were drawn to scan through each embryo, and constricting cells sitting on the left and right boundary of the constriction domain for each sampling line were marked with red dots. (D) Comparison of the number of constricting cells in WT and *fwd* RNAi embryos. Note that the average number of the constricting cells was smaller than 12 due to the sampling method we use. Error bar stands for s.d.; n.s.: not significant. Two tailed, unpaired Student's t test.

**(E)** Cross section view showing basal myosin change over time. Myosin is labeled with Sqh-mCherry. Green dotted line outlines the base of ventral furrow based on E-cadherin-GFP signal. Scale bars, 20 µm. N=4 embryos for both wildtype and *fwd* RNAi embryos.

### Supplementary Figure 4

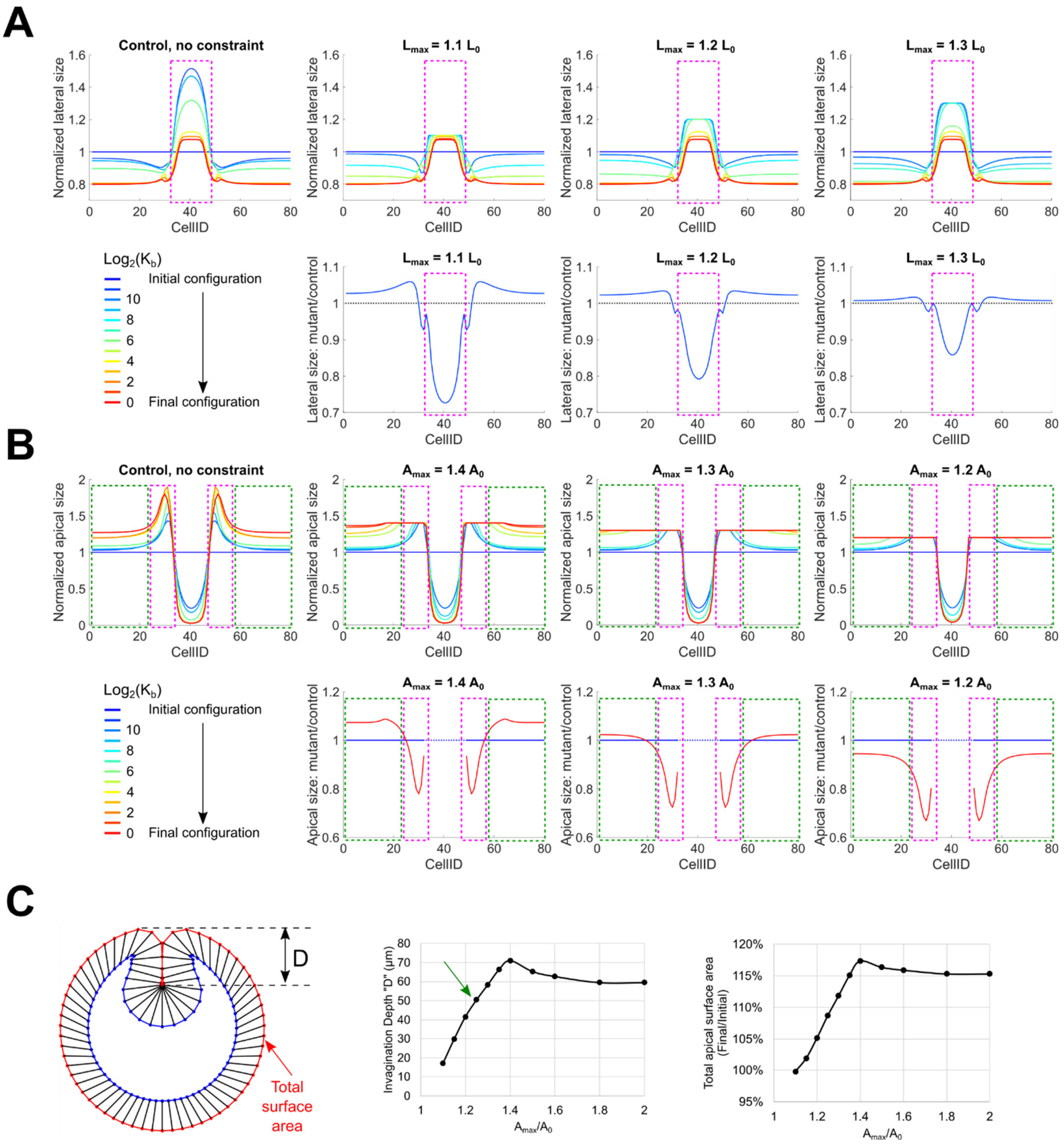

**Supplementary Figure 4. The impact of restricting lateral expansion and apical expansion on different cell populations.**

**(A)** The impact of restricting lateral expansion on cells located at different dorsal-ventral positions. In wildtype situation, only the cells within the constriction domain undergo cell lengthening (magenta box). Therefore, only these cells are substantially influenced by the restriction on lateral expansion.

**(B)** The impact of restricting apical expansion on cells located at different dorsal-ventral positions. In wildtype situation, the flanking cells adjacent to the constriction domain expand the most (up to 2-fold, magenta box). The remaining ectodermal cells also expand their apical domain moderately during the folding process (up to ~1.3-fold, green box). When $A_{max}/A_0 > 1.3$, only the flanking cells are subjected to the constraint. When $A_{max}/A_0 < 1.3$, both the flanking cells and the ectodermal cells are subjected to the constraint.

**(C)** The impact of restricting apical expansion on invagination depth. Left: measurement of invagination depth "D" and total apical area at the end of invagination. Middle and right: "D" and total apical area as a function of $A_{max}/A_0$. When $A_{max}/A_0 > 1.3$, final invagination depth and total apical area are comparable to the control. When $A_{max}/A_0 < 1.3$, final invagination depth and total apical area are negatively impacted, and the level of impact is strongly associated with $A_{max}/A_0$.
